# Overwhelming presence of non-food vegetation undermines the reliability of NDVI as a surrogate of forage abundance for a large herbivore in a tropical forest habitat

**DOI:** 10.1101/356311

**Authors:** Hansraj Gautam, Evangeline Arulmalar, Mihir R Kulkami, TNC Vidya

## Abstract

The use of remotely-sensed vegetation indices has increased in wildlife studies but field-based support for their utility as a measure of forage availability largely comes from open-canopy habitats. In this paper, we assessed whether the popular vegetation index, NDVI, actually represents forage availability for Asian elephants in a southern Indian tropical forest. We found that the number of food species was a small percentage of all plant species, and the abundance of food species compared to total species abundance varied across different vegetation categories. NDVI was not a good measure of food abundance in any vegetation category partly because of a) small to moderate proportional abundances of food species relative to the total abundance of all species in that category (herb and shrub categories), b) abundant overstorey vegetation resulting in low correlations between NDVI and food abundance despite a high proportional abundance of food species and a concordance between total abundance and food species abundance (graminoid category), and c) the relevant variables measured and important as food at the ground level (count and GBH) not being related to primary productivity (trees and recruits). NDVI had a negative relationship with the total abundance of graminoids, which represent a bulk of elephant and other herbivore diet, because of the presence of other vegetation types and canopy cover that positively explained NDVI. We also found that spatially interpolated total graminoid abundance modelled from field data outperformed NDVI in predicting total graminoid abundance, although interpolation models of food graminoid species abundance were not satisfactory. Our results reject the utility of NDVI as a surrogate of elephant forage abundance in tropical forests with multistorey vegetation, a finding that has implications for studies of other herbivores in such habitats.

## Introduction

The distribution and abundance of food resources are among the principal habitat descriptors affecting foraging, movement, habitat selection, and distribution of animals (Pyke *et al.* 1977 Johnson *et al.* 2001, van Beest *et al.* 2010). Food abundance is especially important in shaping the ecology of large herbivores because allometrically decreasing digestive efficiency and increasing dietary requirement with increasing body size has ramifications on foraging time and ranging (Hanley 1982, Demment 1982, Owen-Smith 1988, du Toit and Yetman 2005, Ofstad *et al.* 2016). Therefore, reliable quantification of food resources is crucial to understanding the ecological relationships of large herbivores with their habitat. However, since large herbivores are usually wide-ranging (Ofstad *et al.* 2016), reliable assessment of forage distribution based on field inventory of food species can be demanding in terms of time, effort, and logistics (see Saïd *et al.* 2005), and ecologists have turned towards remote sensing tools for forage distribution and habitat assessment (see Leyequien *et al.* 2007, Pettorelli *et al.* 2011, He *et al.* 2015). Vegetation indices derived from satellite-based remote-sensing data, such as Normalized Difference Vegetation Index (NDVI, Rouse *et al.* 1974), are increasingly being used as indirect measures of forage abundance and habitat quality for developing species distribution or habitat suitability models, and in studies of habitat selection, animal movement and ranging, and population ecology (reviewed in Pettorelli *et al.* 2005, 2011, Leyequien *et al.* 2007, He *et al.* 2015). Animal-habitat relations have been analysed using NDVI in wide-ranging animals such as African buffalo (e.g. Ryan *et al.* 2012), other ungulates (e.g. Pettorelli *et al.* 2006, Hamel *et al.* 2009, Prokopenko *et al.* 2017), marsupials (Youngentob *et al.* 2015), primates (e.g. Zinner *et al.* 2001, Willems *et al.*2009), and elephants (e.g. Murwira and Skidmore 2005, Young *et al*. 2009, Marshal *et al.* 2010, Rood *et al.* 2010, Bohrer *et al.* 2014). However, while NDVI has been found to be a good surrogate of total vegetation biomass/primary productivity in various habitats (savannahs: e.g. Sjöström *et al.* 2009, Wu *et al.* 2013; savannah-steppe mixed landscapes: e.g. Sannier *et al.* 2002; shrublands: e.g. Wilson *et al.* 2011; tropical forests: e.g. Roy and Ravan 1996, Madugundu *et al.* 2008, Das and Singh 2016), tests of whether NDVI or other similar indices actually reflect the abundance or quality of vegetation relevant to the focal animal are unfortunately rare and largely restricted to open habitats such as grasslands and savannahs (e.g. Kawamura *et al.* 2005, Ryan *et al.* 2012, Zengeya *et al.* 2013). We could find only a single such study in temperate forest (Borowik *et al.* 2013) and one from a tropical forest habitat (Willems *et al*. 2009), despite the use of NDVI in studies of animal ecology in such habitats (Zinner *et al.* 2001, Rood *et al.* 2010, Srinivasaiah *et al.* 2012, Marasinghe *et al.* 2015, Lakshminarayan *et al.* 2015, Rahman *et al.* 2017). If remotely sensed indices do not actually reflect forage abundance, their use in studies of foraging and habitat use would lead to artefactual results. We, therefore, compared NDVI with detailed vegetation inventory data collected in the field. Our findings highlight the limitations of NDVI in reliable quantification of forage abundance for a large herbivore, the Asian elephant, in a tropical forest in southern India.

The possible limitations of using remotely-sensed indices to assess herbivore forage in forest habitats stem from the following: **1**) the complexity of multi-storey vegetation structure in forest habitats, and **2**) the small proportion of all plant species in forests that the focal herbivore feeds upon. The first limitation arises from the fact that satellite-based vegetation indices assess greenness of the vegetation detected from above, implying that more abundant productive strata (e.g. tree canopy) will contribute more to such indices than the understorey. Therefore, while NDVI may be correlated with vegetation accessible to herbivores in grasslands, savannahs, open woodlands, and other open habitats with sparse tree cover, it may reflect largely inaccessible vegetation in forests, in which the top tree canopy layer is more closed and constitutes the bulk of the primary productivity detected from the satellite. Therefore, what NDVI detects may not be relevant to foraging by herbivores (see Borowik *et al.* 2013). The second limitation is dependent on the extent of forage selectivity by the herbivore and may be greater in forests than in open areas because of the diversity of species available. The correlation of NDVI with accessible vegetation may suffice in mapping food resources of generalist herbivores (e.g. moose, Belovsky 1978, and African savannah elephants, Owen-Smith 1988) that feed on most components of the accessible vegetation. However, quantification of food resources for selective foragers (e.g. primarily frugivorous primates, see Wheeler *et al.* 2013; browsers like kudu, Owen-Smith and Novellie 1982, giraffe, Pellew 1984, reindeer and musk-oxen, Kazmin *et al.* 2011; and grazers like rhinoceros and hippopotamus, Owen-Smith 1988) would be more complex if their food species represent only a fraction of the accessible vegetation in the habitat.

Elephants are mega-herbivores that feed on a wide variety of plants (McKay 1973, Owen-Smith 1988, Sukumar 1990, Baskaran *et al.* 2010) and have often been classified as generalist feeders (Owen-Smith 1988, Sukumar 1990, but see Owen-Smith and Chafota 2012). However, despite their wide dietary niche compared to those of other sympatric herbivores (Ahrestani *et al.* 2012) and wide distribution across south Asia in a variety of habitats (Hedges *et al*. 2008), Asian elephants may exert considerable choice during feeding in any particular season/habitat, resulting in a small number of plant species forming a large proportion of their diet (Sukumar 1990, Easa 1999, Baskaran *et al.* 2010). The bulk of the diet of Asian elephants is represented by grasses (McKay 1973, Easa 1999, Baskaran *et al*. 2010) despite their lower availability (in terms of biomass) compared to browse (trees, shrubs and non-grass herbaceous plants) in forests. The rich plant diversity of the tropical forests that Asian elephants inhabit (see Myers 2000, Hedges *et al.* 2008) may further lower the proportion of elephant food species there compared to the proportion of herbivore food species in less diverse or grass-rich savannah habitats. Therefore, the notion of generalist/selective foraging does not directly map on to dietary niche breadth, and Asian elephants may be considered selective feeders in the context of the proportion of all plant species available in their habitat that they feed upon. The small proportion of food plants, along with access to only the lower strata of vegetation (bark and stems of trees and shrubs in addition to grass), may result in remotely-sensed indices that measure overall vegetation productivity not adequately capturing forage availability. Therefore, to assess whether NDVI can be used as a proxy for forage availability in a tropical forest, we set out to address the following questions:

1. Does the seasonal and spatial variation in elephant food species abundance match the distribution of total species abundance in different vegetation categories (strata) in a tropical forest? If the abundance of food plants in a vegetation category is not in tight synchrony with overall vegetation abundance in that category, NDVI will be unlikely to capture variations in food plant availability even if it reflects total abundance.
2. How well does NDVI capture the fine scale distribution of a) total species abundance and b) food species abundance in different vegetation categories? The first part will tell us whether NDVI can be used to measure overall (within a vegetation category) plant abundance, especially in the graminoid understorey that is important for elephants and other large herbivores. The second part will assess how well NDVI can map the actual abundance of food species, given the presence of non-food vegetation, in addition to the inaccessibility of vegetation categories that NDVI might measure. We found that NDVI was negatively correlated with graminoid abundance. Therefore, we also investigated c) how the relationship between graminoid abundance and NDVI is linked to the abundance of non-graminoid vegetation.
3. Can a spatial interpolation model be used as an alternative to NDVI as a reliable spatial model of graminoid abundance? Since mapping the spatial distribution of graminoids would be useful in understanding the ecology of various herbivores, we wanted to find out if spatial interpolation from ground data would be as useful as or more useful than NDVI in carrying out such mapping.

Since the Asian elephant has a wide dietary niche breadth, it is expected that the correlation between NDVI and forage abundance would be higher in this species than in other herbivores. Therefore, if NDVI is not a good surrogate of forage abundance in this species, it is likely to be less useful in other, more selective feeders.

## Material and Methods

### Study area

The study was carried out from September 2011 to May 2012 in Nagarahole National Park (area: 644 km^2^, 11.85304°-12.26089° N, 76.00075°-76.27996° E), which lies in the Nilgiris-Eastern Ghats landscape in southern India (Figure 1a). Nagarahole National Park is contiguous with the forests of Bandipur Tiger Reserve, Madikeri Forest Division, and Wayanad Wildlife Sanctuary, which together offer a large contiguous forest landscape with a variety of habitat types to the wide-ranging Asian elephant. The study area can be divided into three broad forest types with multistorey vegetation structure: moist deciduous forest, dry deciduous forest, and teak plantation (Pascal 1982). Kabini and Nagarahole are small, perennial rivers that run through Nagarahole National Park. Between Nagarahole National Park and Bandipur Tiger Reserve lies the Kabini reservoir, which is also perennial, but with widely fluctuating water levels depending upon the season. There are also many small waterholes in the park. The park is home to a variety of large and small herbivores and carnivores. Elephant density in Nagarahole National Park has been estimated at1.74/sq. km through dung sampling methods (AERCC 2006).

**Figure 1.**
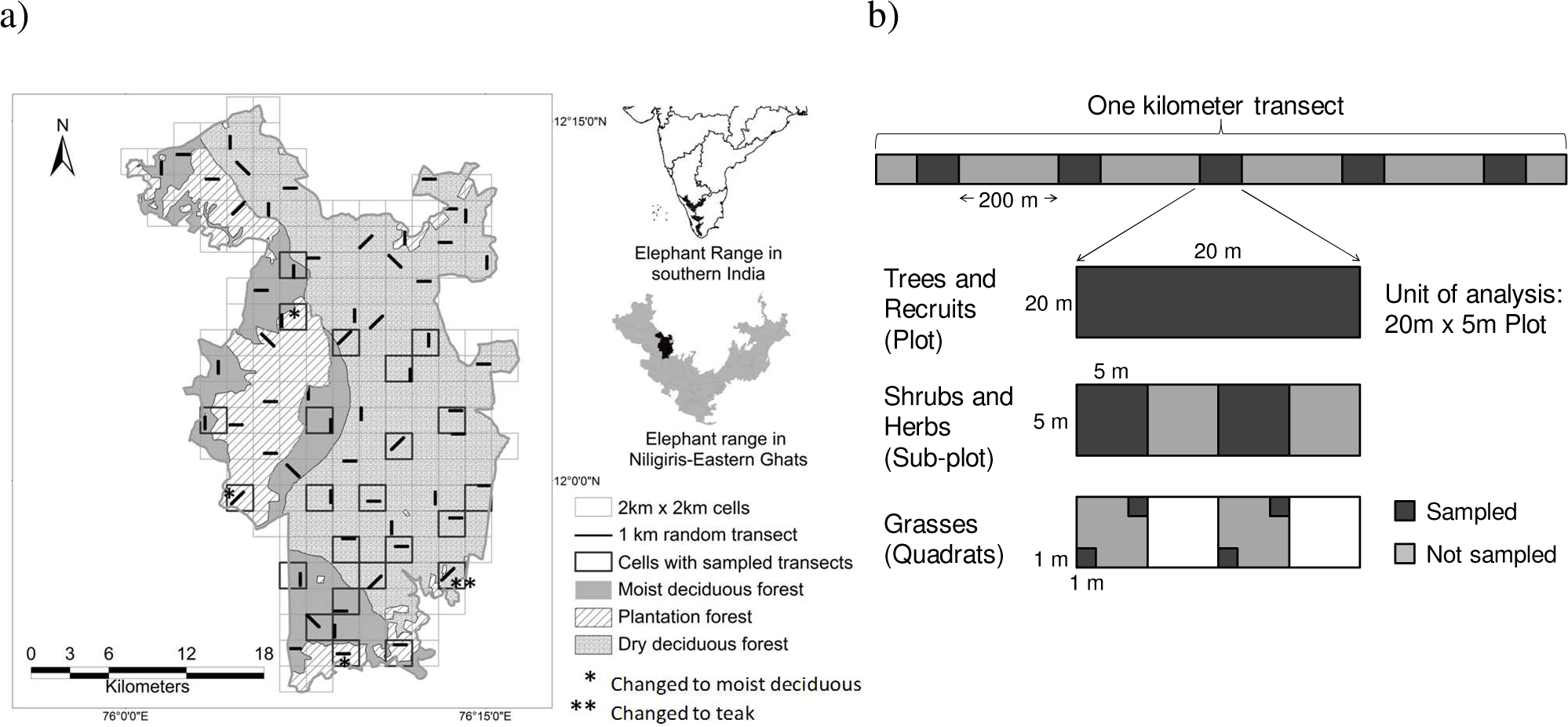
a) Map showing the study area and the location of 60 randomly placed 1-km transects within different forest types (based on Pascal 1982) in Nagarahole National Park. Sampled transects are marked with dark (2km × 2km) cell boundaries. Transects in reassigned forest types are marked with asterisks. b) Schematic of the sampling regime showing a 1-km line transect and the vegetation plots within that were sampled for different plant categories.

### Vegetation Sampling

We gridded Nagarahole National Park into 2 km × 2 km cells, chose randomly selected cells, and placed 1-km long line-transects according to a stratified sampling regime (Figure 1a). Due to logistics and time constraints, the northern parts of the park were not sampled and we sampled 17 transects during the wet season (17 September - 13 December 2011) and 22 (of which 16 had been sampled in wet season) transects during the following dry season (29 February - 14 May 2012). Transects were selected in order to sample the three forest types proportionate to the areas they covered. Ground-truthing revealed a smaller area covered by teak plantations compared to that based on Pascal’s (1982) map, resulting in reassigning three “teak plantation” transects as moist deciduous forest and one “dry deciduous” transect as teak plantation transect (shown by asterisks in Figure 1a). Therefore, 9, 6, and 2 transects were sampled in the dry deciduous forest, moist deciduous forest, and teak plantation, respectively, in the wet season, and 11, 9, and 2, respectively, in the dry season.

Along each transect, we sampled five plots of 20 m × 5 m, spaced at 200 m intervals, and collected abundance data from different vegetation categories: trees, recruits, shrubs, herbs, and graminoids. Graminoids included mostly Poaceae members but also a few Cyperaceae and Juncaceae species, and herbs included non-graminoid herbaceous plants. Individuals of tree species with girth at breast height (GBH) of at least 10 cm were classified as trees, while individuals of these species with a girth of less than 10 cm were classified as recruits. We recorded the number of individuals and GBH of trees and the number of individuals of recruits within the 20m × 5m plots. Within each 20 m × 5 m plot, we sampled two 5 m × 5 m sub-plots for shrubs and herbs, and four 1 m × 1 m quadrats for graminoids (Figure 1b). Shrub, herb, and graminoid abundance was assessed by visually estimating the percentage cover of each species within the sampling sub-plot / quadrat (Gautam *et al.* 2017; visual estimation has also been used successfully by Tsalyuk *et al.* 2017). We photographed the canopy (using a Canon SX120 IS digital camera - 10 megapixels) from a height of 1 m at each of the 1m × 1m quadrats in order to measure canopy cover. We sampled 85 tree plots, 170 shrub/herb sub-plots, and 340 graminoid quadrats from 17 one-km line transects during the wet season of 2011, and 110 tree plots, 220 shrub/herb sub-plots, and 440 graminoid quadrats during the dry season of 2012. Identification of plant species was carried out by two taxonomy experts: Dr. Ravikumar (FRLHT, Bangalore) identified the herb, shrub and tree species, and Dr. Girish Potdar (Yeswantrao Chavan College of Science, Karad) identified the graminoid species.

### Processing of data for vegetation and elephant food species abundance

We used the sum of species abundance of all species in each vegetation category as a measure of total species abundance for that category. A strong relationship had been previously found in our study area between percentage cover and fresh biomass, both for individual species and for sum of species abundance (Gautam *et al.* 2017). We used percentage covers of species (of the same vegetation category) to obtain the sum of species abundance of graminoids, herbs, and shrubs. Counts of individuals (for trees and recruits) and girth (for trees) were added across species to obtain different measures of the sum of species abundance of trees and recruits. The sum of elephant food species abundance was calculated separately to obtain elephant food species abundance in each vegetation category. The list of elephant food species in the study area (see Supplementary material Appendix 1 Table 1) was compiled from 1) our visual observations (during 2012-2013) of feeding by elephants and debarking marks on trees in the study area, and **2**) from previous studies of elephant feeding ecology in neighbouring areas: Mudumalai Tiger Reserve (Sivaganesan 1991), Wayanad Wildlife Sanctuary (Easa 1999), and parts of the Nilgiri Biosphere Reserve (Baskaran *et al* 2010).

**Table 1.** Number of 1 km line-transects sampled and used in ANOVAs on the abundance of elephant food species and all species. Each transect has five plots.

| Analysis | Vegetation category | Dry deciduous transects |  | Moist deciduous transects |  | Teak plantation transects |  |
| --- | --- | --- | --- | --- | --- | --- | --- |
|  |  | Sampled | Used | Sampled | Used | Sampled | Used |
| Nested, repeated measures ANOVA | Graminoids, herbs, shrubs, recruits (wet season) | 9 | 6 | 6 | 6 | 2 | NA |
|  | Graminoids, herbs, shrubs, recruits (dry season) | 11 | 6 | 9 | 6 | 2 | NA |
| Nested ANOVA | Trees (count and GBH) | 11 | 9 | 9 | 9 | 2 | NA |

\* For moist deciduous forest, the same six transects that were sampled in the wet season were considered in the dry season for the analysis and the additional three transects sampled in the dry season were excluded. In case of dry deciduous forest, of the nine transects sampled in the wet season, we excluded a transect at the edge of the protected-area boundary, one that was at the transition between dry deciduous and teak plantation, and another one from two transects that were very close to each other in order to achieve a balanced design.

Photographs taken of the canopy were processed in Black Spot Leaf Area Calculator (Varma and Osuri 2013) to obtain canopy cover by classifying the image into dark (regions that intercept light) and light areas based on digital values of the image pixels. The canopy covers above the four 1m × 1m quadrats in each 20m × 5m plot were averaged and used as the representative value for that plot.

### Satellite data processing

High resolution satellite images, chosen for least cloud cover over the study area, from the middle of the wet season (16/11/2011, LISS 3, 23.5m resolution) and the dry season (03/03/2012, LISS 4, 5.8m resolution) were obtained from National Remote Sensing Centre, Hyderabad, India, and processed according to standard image processing procedure (Lillesand *et al.* 2004). Radiometric correction by dark-pixel subtraction and resampling (Lillesand *et al.* 2004) to a common (23.5m) resolution (resampling was done for the LISS 4 image which was of 5.8m resolution) was performed. NDVI was calculated from near infra red (NIR) and red (R) bands by the formula, NDVI=(NIR-R)/(NIR+R) (Rouse *et al.* 1974). Satellite image processing and NDVI calculation was carried out in Erdas Imagine 9.1(Leica Goesystems 2005) and NDVI values for each plot were extracted using Spatial Analyst of ArcMap 10.1(ESRI 2012).

### Data analysis

**1. Seasonal and spatial variation in elephant food species abundance and total species abundance in different vegetation categories**

We used nested ANOVAs (Doncaster and Davy 2007, pp. 214-216) to examine spatial and seasonal variation in elephant food species abundance and total species abundance. Nested repeated-measures ANOVAs were carried out on elephant food species or total species abundance, separately for each vegetation category except trees, with season as a within-subject effect, plot (20m × 5m, random subject) nested under transect (random effect), and transect nested under forest type (fixed effect). The nested design was used to account for variations arising from different spatial scales i.e., the forest type and transect scale. Since equal numbers of transects in each forest type were required across seasons to carry out nested ANOVAs, 6 transects each from the dry deciduous and moist deciduous forests that were sampled in both seasons were used, and the remaining transects were excluded (Table 1. Since the count or girth of trees was not expected to change appreciably across a single season, nested ANOVAs, with plots nested under transects and transects nested under forest type, were carried out for elephant food tree species and all tree species separately. Nine transects each from dry and moist deciduous forests sampled during the dry season were used for these ANOVAs (Table 1). We regressed elephant food species abundance on the total species abundance, separately for each vegetation category and season, to examine the extent to which elephant food species abundance was predicted by total species abundance. Forest type was used as a categorical predictor in all regressions. Data from all transects were used for these regressions.

**2. Relationship of NDVI with total species abundance and elephant food species abundance**

In order to understand the extent to which NDVI explains total species and elephant food species abundance, we carried out separate general linear regressions of the two on NDVI, using forest type as a categorical predictor. These regressions were carried out for each vegetation category and season. We found, based on the analysis above, that NDVI was significantly negatively correlated with total and elephant food graminoid abundance in the dry season and with total graminoid abundance in wet season (see Results). Since graminoids are important in elephant and other ungulate diet, we examined whether their negative relationship with NDVI was linked to different non-graminoid vegetation variables, which might have inhibitory effects of shade on graminoid abundance. In order to do this, we looked at the vegetation variables that contributed most to NDVI by performing best-subsets regression of NDVI (dependent variable) on continuous predictors that included the total abundance of each vegetation category and canopy cover, along with forest type as a categorical predictor. These analyses were performed separately for the wet and dry seasons. We then performed best-subsets regression of total graminoid species abundance on other vegetation variables (total abundance of trees, recruits, shrubs, herbs) and canopy cover to find out whether non-graminoid vegetation variables were affecting graminoid abundance. Best subsets regression models were selected using Mallows’ *Cp* statistic (Mallows 1973; best models have lowest *Cp* values closest to the number of parameters, p). Effect sizes of predictor variables in the multiple regression models were estimated using η^2^ (SS_effect_/SS_corrected total_, see Fritz *et al.* **2012**) which was calculated from tables of univariate tests of significance.

**3. Comparing NDVI versus spatial interpolation in mapping graminoid abundance**

The use of NDVI to model graminoid abundance was compared with the spatial interpolation method in which graminoid abundance data collected from the field were interpolated in Spatial Analyst of ArcMap 10 using ordinary kriging. Of the 110 plots sampled in the dry season, 55 randomly selected plots were used as the training dataset for preparing an interpolated spatial model, whereas 50plots were used to test how well interpolation could predict the sum of graminoid abundance (5 plots were excluded because they had burnt undergrowth). Spatial interpolation was repeated 10 times in the manner above, selecting 55 random plots as the training dataset each time (however, in every training dataset, 4 plots with maximum northing, southing, easting and westing were not random, in order to retain the maximum spatial extent of sampling). This analysis was also done on data from the wet season with45 plots in the training dataset and 40 in the verification dataset. Coefficients of determination (*R*^2^) were obtained after regressing the observed graminoid abundance on the modelled graminoid abundance (from the 10 datasets). Using the same verification plots from the 10 datasets, the observed graminoid abundance was also regressed on NDVI to obtain R^2^ for the NDVI model. Wilcoxon matched-pairs test was used to compare *R*^2^ values between the interpolation model and the NDVI model. Similarly, kriging and NDVI models were also compared using data on abundance of only the elephant food grass species. Whether increasing the sample size of the training dataset from 55 to 75 plots (30 verification plots) improved the prediction from interpolation was also examined using the dry season dataset (insufficient data from the wet season) on total grass abundance and 10 iterations.

Statistica 7 software (StatSoft 2007) was used to carry out all statistical tests.

## Results

**1. Seasonal and spatial variation in elephant food species abundance and total species abundance in different vegetation strata**

We recorded 435 plant species from vegetation sampling, which included 77 graminoid species, 181 herbaceous (herbs, creepers and climbers) species, 48 shrubs, 120 recruits, and 79 trees. Species sometimes overlapped across vegetation categories (such as trees and recruits, and shrubs with GBH>10cm and trees). Sixty of the 435 species (13.79%) were elephant food species (see Supplementary material Appendix 1 Table 1). The mean proportional abundance of food species (elephant food species abundance divided by the total species abundance, and averaged across all plots) was high (~0.8) among graminoids, moderately high (>0.5) for trees and recruits, and low (<0.25) among herbs and shrubs (Table 2).

**Table 2.** Total number of species, number of elephant food species, and mean proportional abundance of food species in different vegetation categories.

| Vegetation category | Total number of species | Number of food species | Mean (SD) proportional food species abundance in the wet season | Mean (SD) proportional food species abundance in the dry season |
| --- | --- | --- | --- | --- |
| Graminoids | 77 | 16 (20.78%) | 0.80 (0.261) | 0.85 (0.266) |
| Herbs | 181 | 9 (4.97%) | 0.23 (0.183) | 0.09 (0.141) |
| Shrubs | 48 | 6 (12.5%) | 0.18 (0.142) | 0.16 (0.136) |
| Recruits | 120 | 28 (23.33%) | 0.58 (0.241) | 0.60 (0.265) |
| Trees | 79 | 25 (31.65%) | NA | 0.54 (0.317; count)<br>0.59 (0.315; GBH) |

There was a fair amount of spatial and seasonal variability in both elephant food species and total species abundance, with 75% (18 out of 24) of the effects (forest type, transect, season, and interactions) being concordantly significant or not between food species and all species (ANOVA tests in Table 3). However, graminoid elephant food species and total species abundance differed in 2 out of 5 effects (Table 3). Forest type significantly affected total abundance in the graminoid and shrub categories, and in the tree category based on counts rather than GBH (Table 3). Graminoids and trees (count) were more abundant in the dry deciduous forest than in the moist deciduous forest, whereas shrubs were more abundant in the moist deciduous forest (see Supplementary material Appendix 2 Table A2 for abundance estimates, Appendix 3 Fig. A1 for plots). Total species abundance in all categories differed significantly across transects (within forest type),except for trees which differed across transects based on tree-count but not based on GBH (Table 3, see Supplementary material Appendix 2 Table A2 for abundance estimates, Supplementary material Appendix 3 Fig. A2 for plots). There was also a significant seasonal effect on total species abundance in all categories examined except for recruits (Table 3), with abundances being lower in the dry season compared to the wet season (Supplementary material Appendix 2 Table A2, Appendix 3 Fig. A1).

**Table 3.**
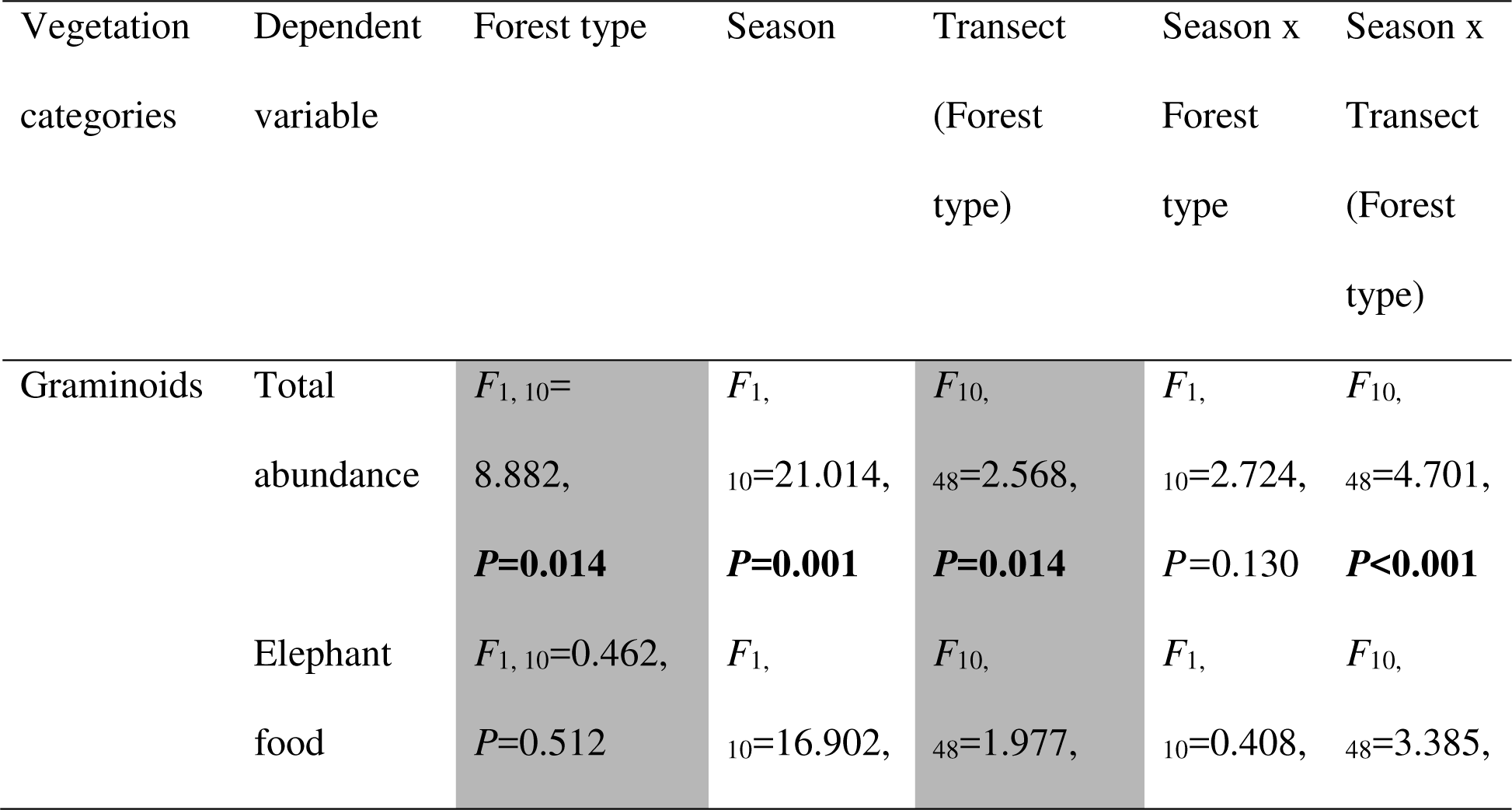
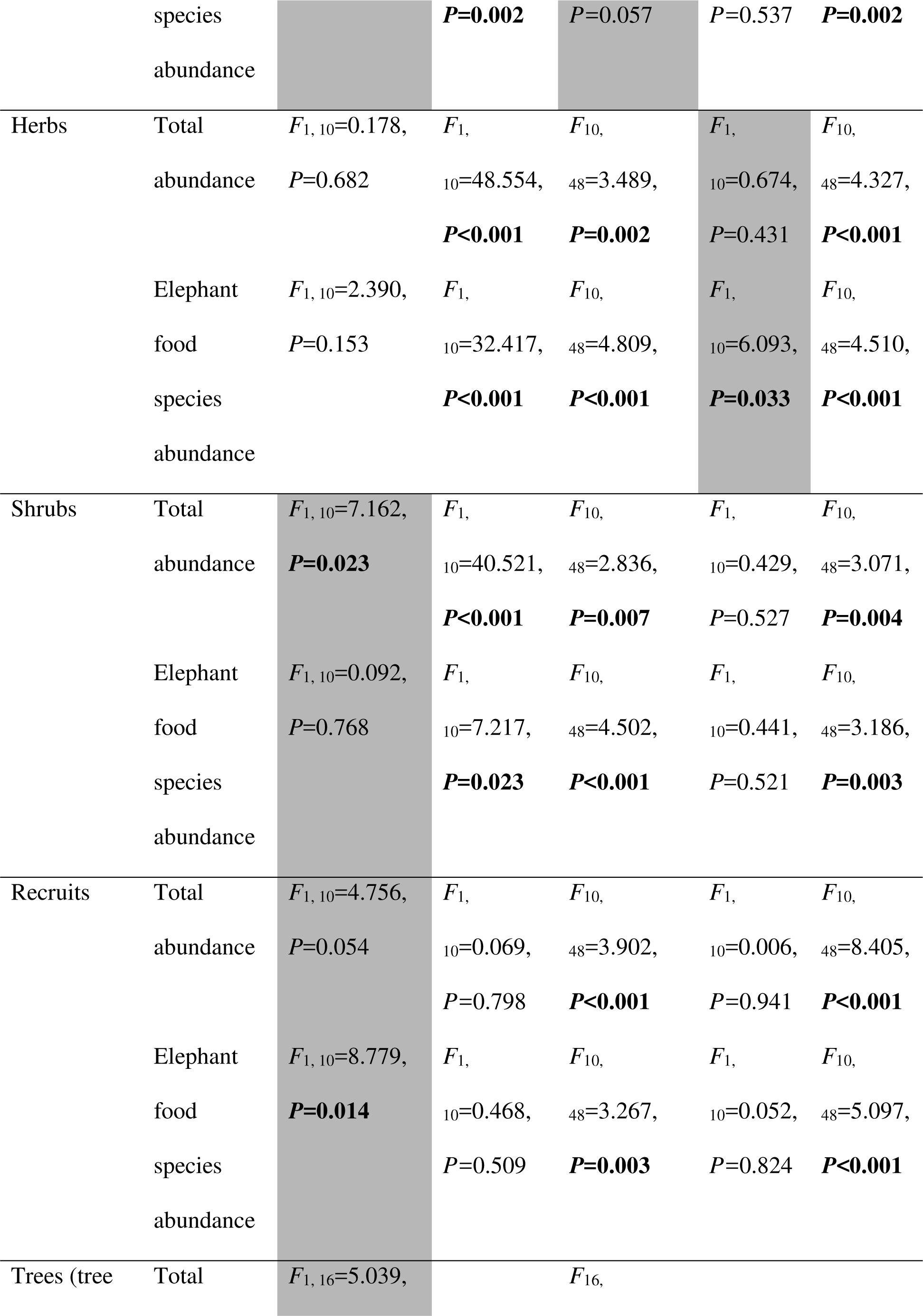
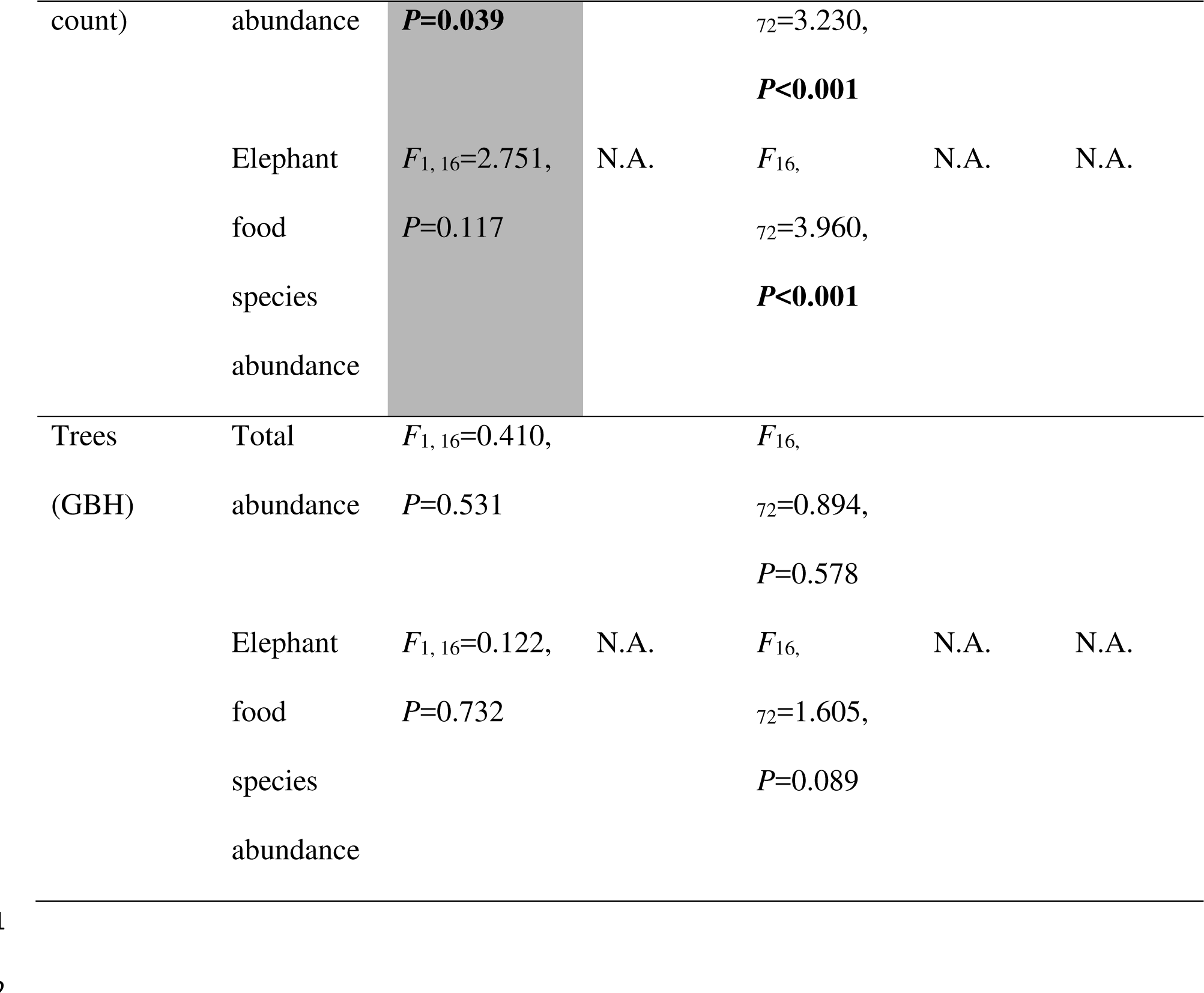
ANOVA results for different vegetation categories showing spatial and seasonal variation in elephant food species abundance and total species abundance. Plot (subject) was nested under transect (random effect), transect nested under forest type (fixed effect) and season was the within-subject repeated measure. Season was not included for tree count and tree GBH. The cells filled with grey point to differences in effects on elephant food species abundance and total species abundance.

Unlike in the case of total abundance, there was no effect of forest type on elephant food species abundance, except for food recruits being more abundant in the dry deciduous forest than in the moist deciduous forest (see Table 3 for ANOVA tests, Supplementary material Appendix 2 Table A2 for abundance estimates, Appendix 3 Fig. A1 for plots). However, elephant food species abundance varied at the level of transects (within forest type) in all categories except for graminoids and tree GBH (Table 3, plots in Supplementary material Appendix 3 Fig. A2). Seasonal differences in elephant food species abundance mirrored those of total species abundance in the respective categories, with food graminoids, herbs, and shrubs, but not food recruit counts, being more abundant in the wet season than in the dry season (Table 3, Supplementary material Appendix 2 Table A2, Supplementary material Appendix 3 Fig. A1).

General regressions showed that total abundance, along with forest type, was significantly related to elephant food species abundance in different categories to varying extents (Table 4). In both the wet and dry seasons, the effect sizes (*η*^2^) of total abundance were large (>0.5) for graminoids, recruits and trees, but low to moderate (less than or close to 0.25) for herbs and shrubs.

**Table 4.**
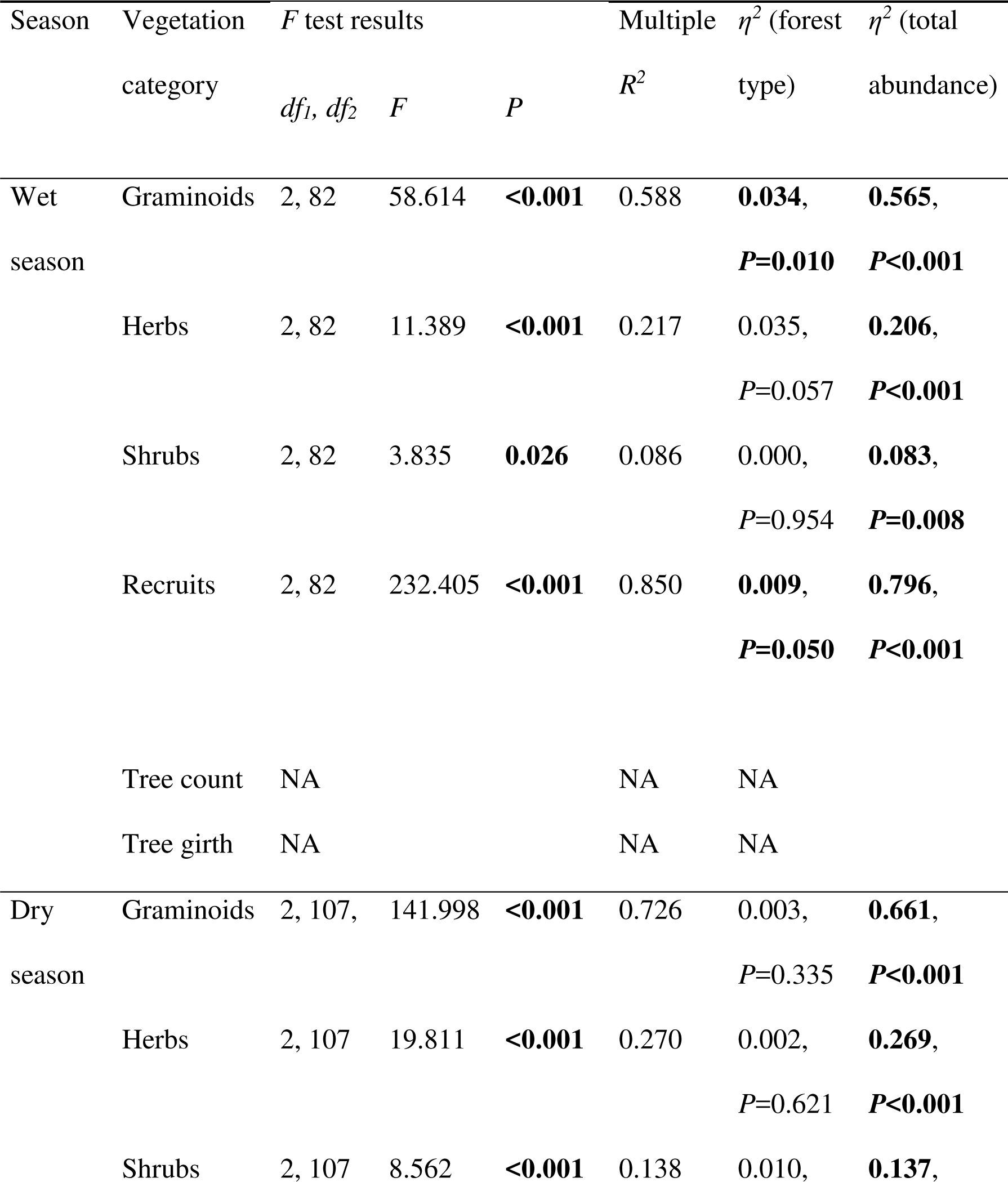
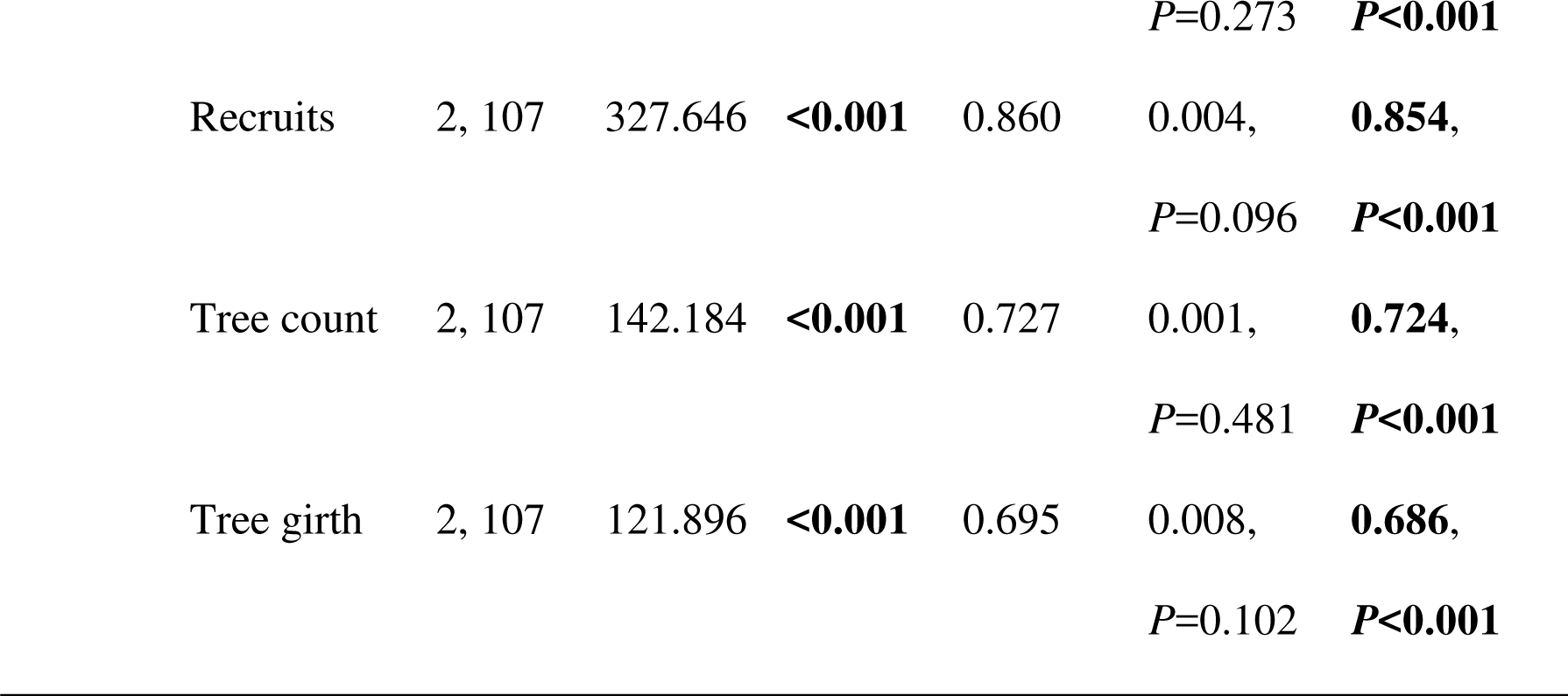
Results of general regressions of elephant food species abundance (dependent variable) on total abundance (continuous predictor) and forest type (categorical predictor) for different vegetation categories. Effect sizes *(η*^*2*^) in this and other tables were calculated based on univariate tests of significance.

| Season | Vegetation category | F test results | | | Multiple $R^2$ | $\eta^2$ (forest type) | $\eta^2$ (total abundance) |
| --- | --- | --- | --- | --- | --- | --- | --- |
| | | $df_1, df_2$ | $F$ | $P$ | | | |
| Wet season | Graminoids | 2, 82 | 58.614 | **<0.001** | 0.588 | **0.034,** | **0.565,** |
|  |  |  |  |  |  | **$P=0.010$** | **$P<0.001$** |
|  | Herbs | 2, 82 | 11.389 | **<0.001** | 0.217 | 0.035, | **0.206,** |
|  |  |  |  |  |  | **$P=0.057$** | **$P<0.001$** |
|  | Shrubs | 2, 82 | 3.835 | **0.026** | 0.086 | 0.000, | **0.083,** |
|  |  |  |  |  |  | **$P=0.954$** | **$P=0.008$** |
|  | Recruits | 2, 82 | 232.405 | **<0.001** | 0.850 | **0.009,** | **0.796,** |
|  |  |  |  |  |  | **$P=0.050$** | **$P<0.001$** |
|  | Tree count | NA |  |  | NA | NA |  |
|  | Tree girth | NA |  |  | NA | NA |  |
| Dry season | Graminoids | 2, 107, | 141.998 | **<0.001** | 0.726 | 0.003, | **0.661,** |
|  |  |  |  |  |  | **$P=0.335$** | **$P<0.001$** |
|  | Herbs | 2, 107 | 19.811 | **<0.001** | 0.270 | 0.002, | **0.269,** |
|  |  |  |  |  |  | **$P=0.621$** | **$P<0.001$** |
|  | Shrubs | 2, 107 | 8.562 | **<0.001** | 0.138 | 0.010, | **0.137,** |

|  |  |  |  |  |  |  |
| --- | --- | --- | --- | --- | --- | --- |
|  |  |  |  |  | *P*=0.273 | *P*<0.001 |
| Recruits | 2, 107 | 327.646 | <0.001 | 0.860 | 0.004, | **0.854,** |
|  |  |  |  |  | *P*=0.096 | *P*<0.001 |
| Tree count | 2, 107 | 142.184 | <0.001 | 0.727 | 0.001, | **0.724,** |
|  |  |  |  |  | *P*=0.481 | *P*<0.001 |
| Tree girth | 2, 107 | 121.896 | <0.001 | 0.695 | 0.008, | **0.686,** |
|  |  |  |  |  | *P*=0.102 | *P*<0.001 |

**2. Relationship of NDVI with total species abundance and elephant food species abundance**

**a) NDVI and total abundance**

General regressions of total abundance on NDVI and forest type for different vegetation categories yielded significant regression models in the case of graminoids and shrubs in the dry and wet seasons (Table 5). However, NDVI, by itself, showed a significant effect only on the total abundance of shrubs in the wet season and total abundance of graminoids in the dry season. The latter effect of NDVI on total graminoid abundance in the dry season was negative, with larger NDVI values indicating lower graminoid abundance.

**Table 5.**
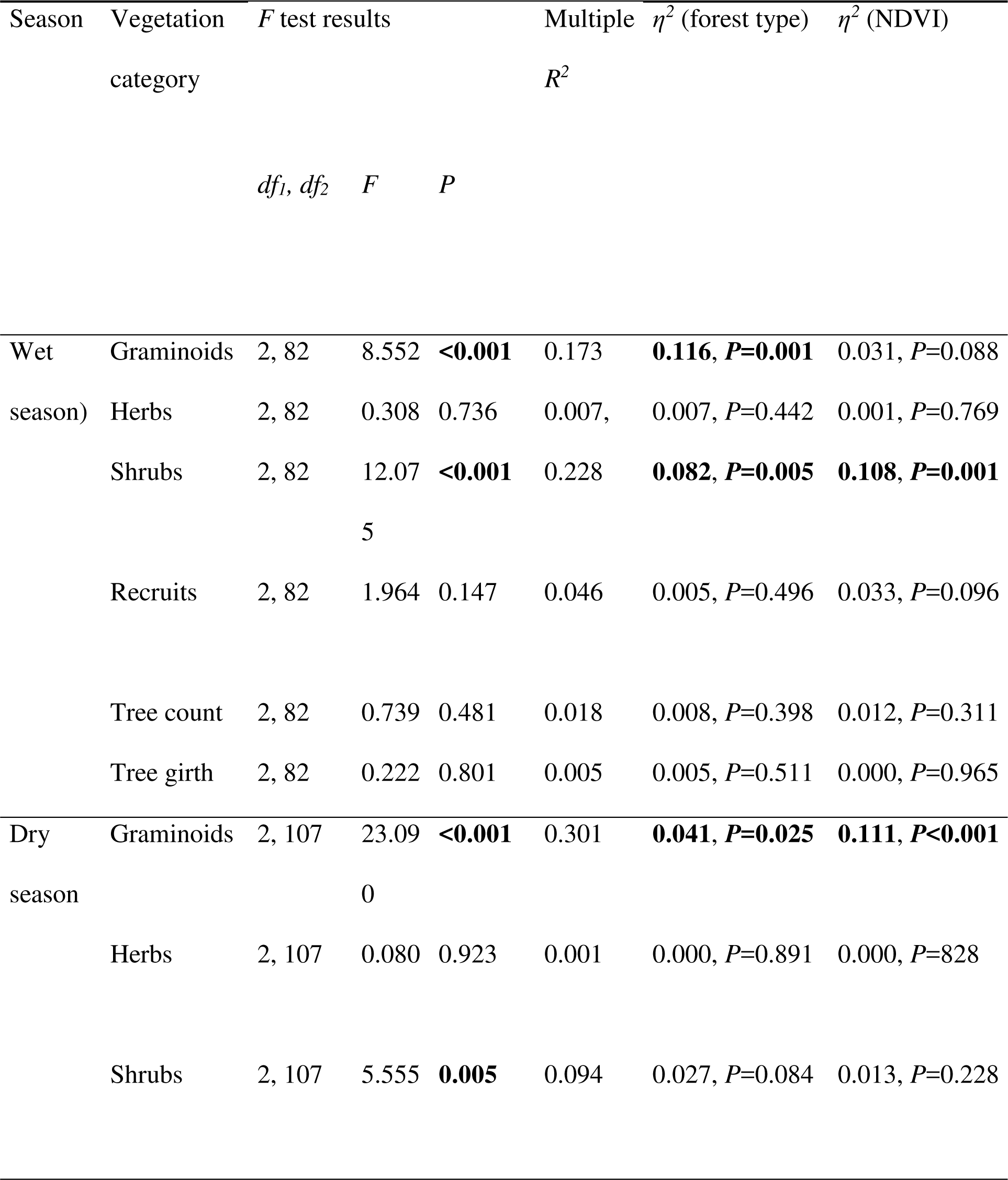
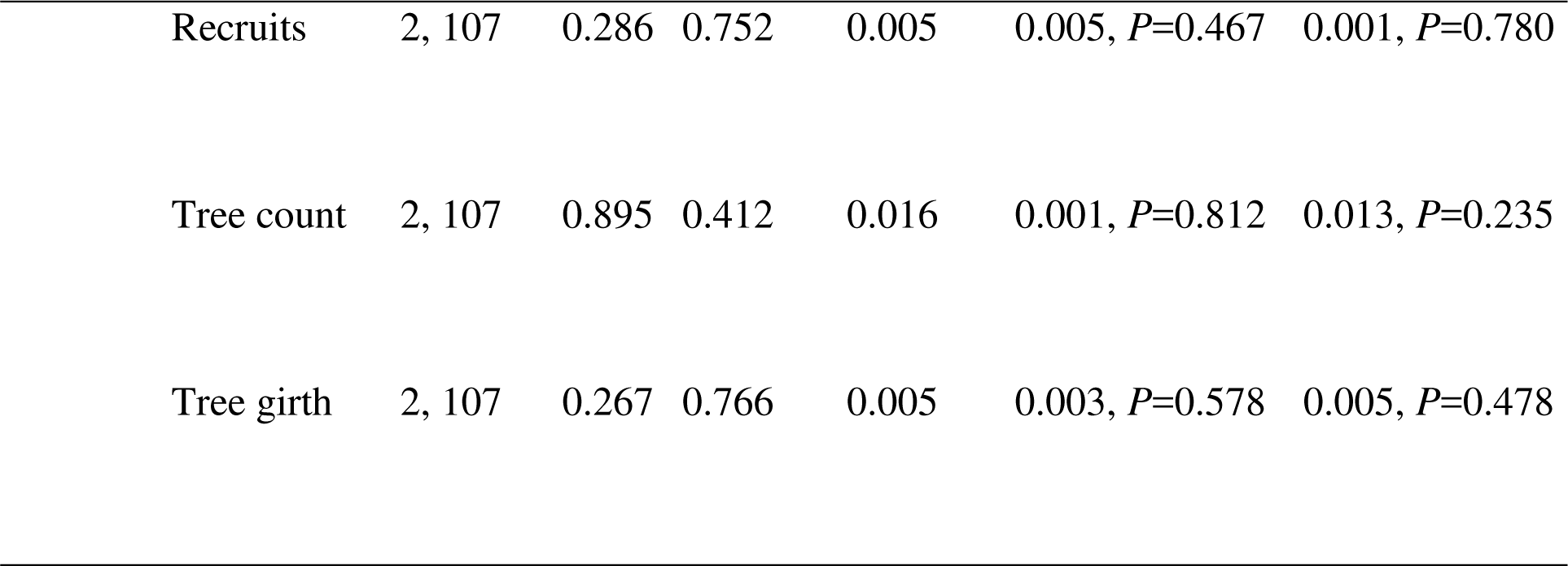
Results of general regressions of total abundance (dependent variable) on NDVI (continuous predictor) and forest type (categorical predictor) for different vegetation categories.

| Season | Vegetation category | *F* test results | | | Multiple $R^2$ | $\eta^2$ (forest type) | $\eta^2$ (NDVI) |
| --- | --- | --- | --- | --- | --- | --- | --- |
|  |  | *df<sub>1</sub>, df<sub>2</sub>* | *F* | *P* |  |  |  |
| Wet season) | Graminoids | 2, 82 | 8.552 | **<0.001** | 0.173 | **0.116, *P*=0.001** | 0.031, *P*=0.088 |
|  | Herbs | 2, 82 | 0.308 | 0.736 | 0.007, | 0.007, *P*=0.442 | 0.001, *P*=0.769 |
|  | Shrubs | 2, 82 | 12.07 | **<0.001** | 0.228 | **0.082, *P*=0.005** | **0.108, *P*=0.001** |
|  |  |  | 5 |  |  |  |  |
|  | Recruits | 2, 82 | 1.964 | 0.147 | 0.046 | 0.005, *P*=0.496 | 0.033, *P*=0.096 |
|  | Tree count | 2, 82 | 0.739 | 0.481 | 0.018 | 0.008, *P*=0.398 | 0.012, *P*=0.311 |
|  | Tree girth | 2, 82 | 0.222 | 0.801 | 0.005 | 0.005, *P*=0.511 | 0.000, *P*=0.965 |
| Dry season | Graminoids | 2, 107 | 23.09 | **<0.001** | 0.301 | **0.041, *P*=0.025** | **0.111, *P*<0.001** |
|  |  |  | 0 |  |  |  |  |
|  | Herbs | 2, 107 | 0.080 | 0.923 | 0.001 | 0.000, *P*=0.891 | 0.000, *P*=828 |
|  | Shrubs | 2, 107 | 5.555 | **0.005** | 0.094 | 0.027, *P*=0.084 | 0.013, *P*=0.228 |

|  |  |  |  |  |  |  |
| --- | --- | --- | --- | --- | --- | --- |
| Recruits | 2, 107 | 0.286 | 0.752 | 0.005 | 0.005, $P=0.467$ | 0.001, $P=0.780$ |
| Tree count | 2, 107 | 0.895 | 0.412 | 0.016 | 0.001, $P=0.812$ | 0.013, $P=0.235$ |
| Tree girth | 2, 107 | 0.267 | 0.766 | 0.005 | 0.003, $P=0.578$ | 0.005, $P=0.478$ |

**b) NDVI and food species abundance**

General regressions of elephant food species abundance in different vegetation categories on NDVI and forest type yielded different regression models in the dry and wet seasons (Table 6). In the wet season, the regressions of NDVI, along with forest type, on food species abundance of herbs and recruits were significant, whereas the regressions were not significant for other vegetation categories. In the dry season, the regressions of food species abundance of graminoids, and trees (tree count) on NDVI and forest type were significant. However, even when the regressions were significant, the effect sizes of NDVI were low (<0.10 in all cases, Table 6).

**Table 6.**
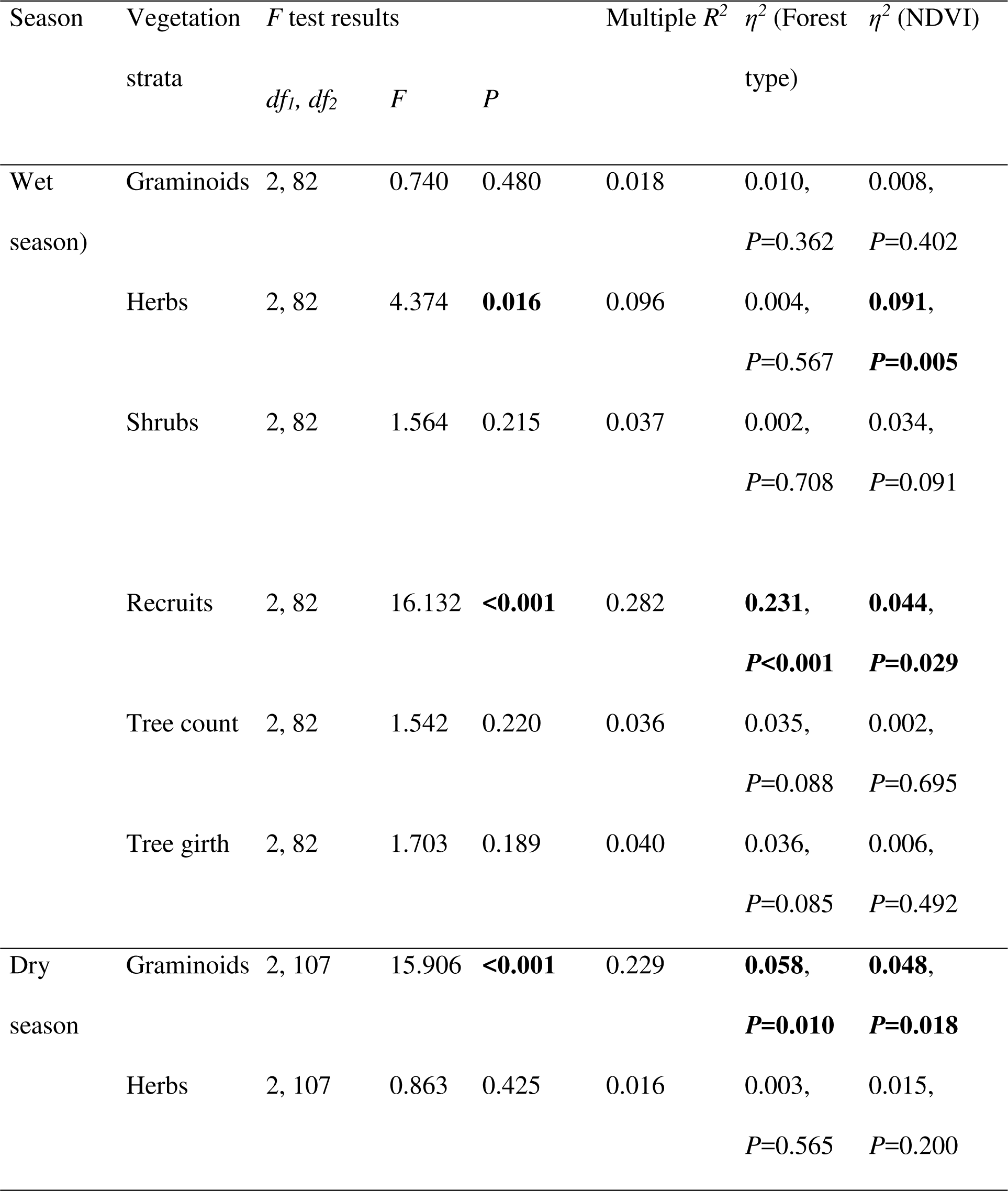
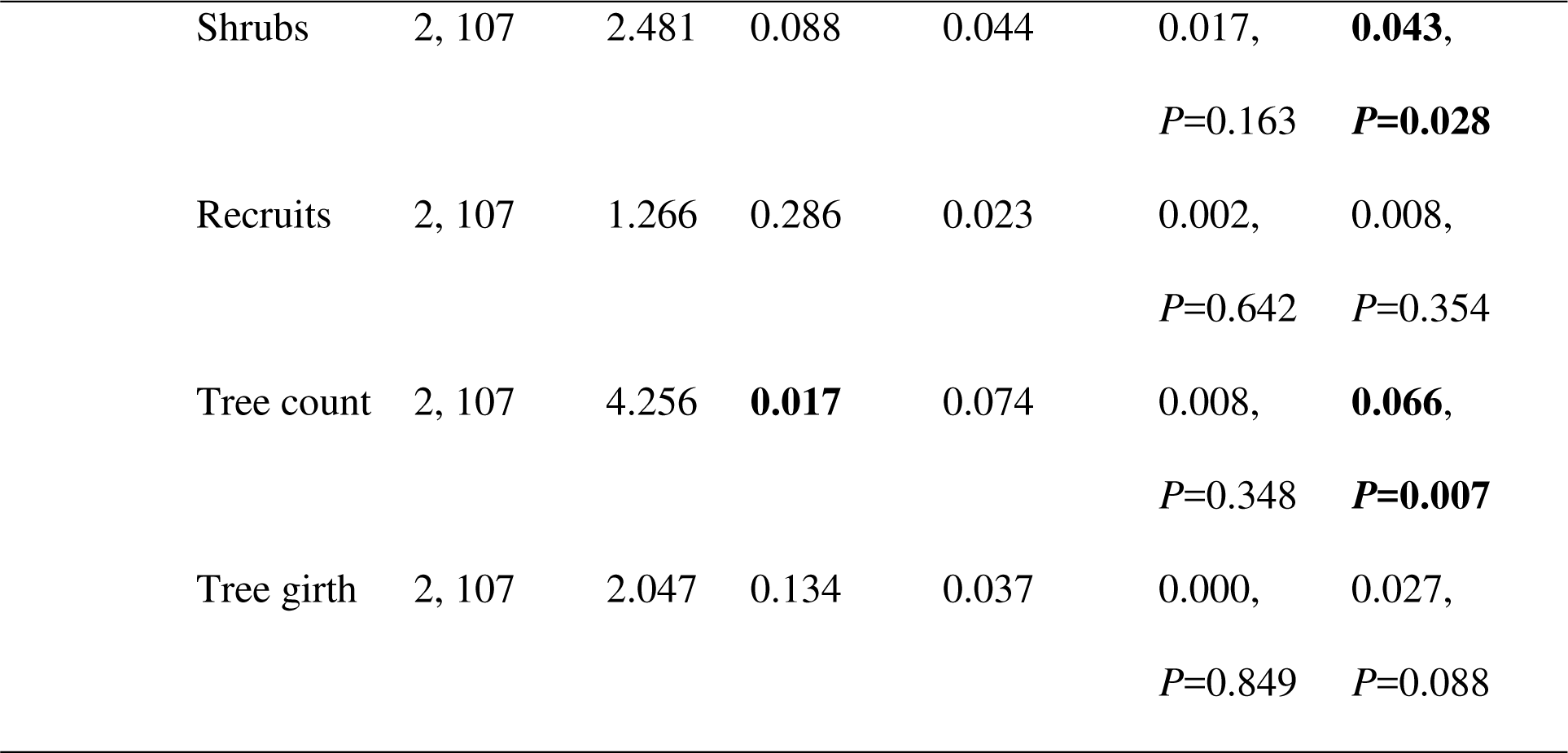
Results of general regressions of elephant food species abundance (dependent variable) on NDVI (continuous predictor) and forest type (categorical predictor) for different vegetation strata.

| Season | Vegetation<br>strata | *F* test results | | | Multiple $R^2$ | $\eta^2$ (Forest<br>type) | $\eta^2$ (NDVI) |
| --- | --- | --- | --- | --- | --- | --- | --- |
| | | $df_1, df_2$ | *F* | *P* | | | |
| Wet<br>season) | Graminoids | 2, 82 | 0.740 | 0.480 | 0.018 | 0.010,<br>*P*=0.362 | 0.008,<br>*P*=0.402 |
|  | Herbs | 2, 82 | 4.374 | **0.016** | 0.096 | 0.004,<br>*P*=0.567 | **0.091**,<br>***P*=0.005** |
|  | Shrubs | 2, 82 | 1.564 | 0.215 | 0.037 | 0.002,<br>*P*=0.708 | 0.034,<br>*P*=0.091 |
|  | Recruits | 2, 82 | 16.132 | **<0.001** | 0.282 | **0.231**,<br>***P*<0.001** | **0.044**,<br>***P*=0.029** |
|  | Tree count | 2, 82 | 1.542 | 0.220 | 0.036 | 0.035,<br>*P*=0.088 | 0.002,<br>*P*=0.695 |
|  | Tree girth | 2, 82 | 1.703 | 0.189 | 0.040 | 0.036,<br>*P*=0.085 | 0.006,<br>*P*=0.492 |
| Dry<br>season | Graminoids | 2, 107 | 15.906 | **<0.001** | 0.229 | **0.058**,<br>***P*=0.010** | **0.048**,<br>***P*=0.018** |
|  | Herbs | 2, 107 | 0.863 | 0.425 | 0.016 | 0.003,<br>*P*=0.565 | 0.015,<br>*P*=0.200 |

|  |  |  |  |  |  |  |
| --- | --- | --- | --- | --- | --- | --- |
| Shrubs | 2, 107 | 2.481 | 0.088 | 0.044 | 0.017, | **0.043,** |
|  |  |  |  |  | *P*=0.163 | ***P*=0.028** |
| Recruits | 2, 107 | 1.266 | 0.286 | 0.023 | 0.002, | 0.008, |
|  |  |  |  |  | *P*=0.642 | *P*=0.354 |
| Tree count | 2, 107 | 4.256 | **0.017** | 0.074 | 0.008, | **0.066,** |
|  |  |  |  |  | *P*=0.348 | ***P*=0.007** |
| Tree girth | 2, 107 | 2.047 | 0.134 | 0.037 | 0.000, | 0.027, |
|  |  |  |  |  | *P*=0.849 | *P*=0.088 |

**c) Relationship between graminoid abundance and NDVI with respect to non-graminoid vegetation abundance**

An examination of the components of vegetation that contributed to NDVI showed that the best regression model of wet season data included the total abundance of shrubs, recruits, and tree count, and explained 20.7% of the variation in NDVI (Table 7). Based on dry season data, the best multiple regression model of NDVI included forest type, canopy cover, and tree count, and explained a much larger variation (63.8%) in NDVI. The top three models for each season are shown in Table 7.

**Table 7.**
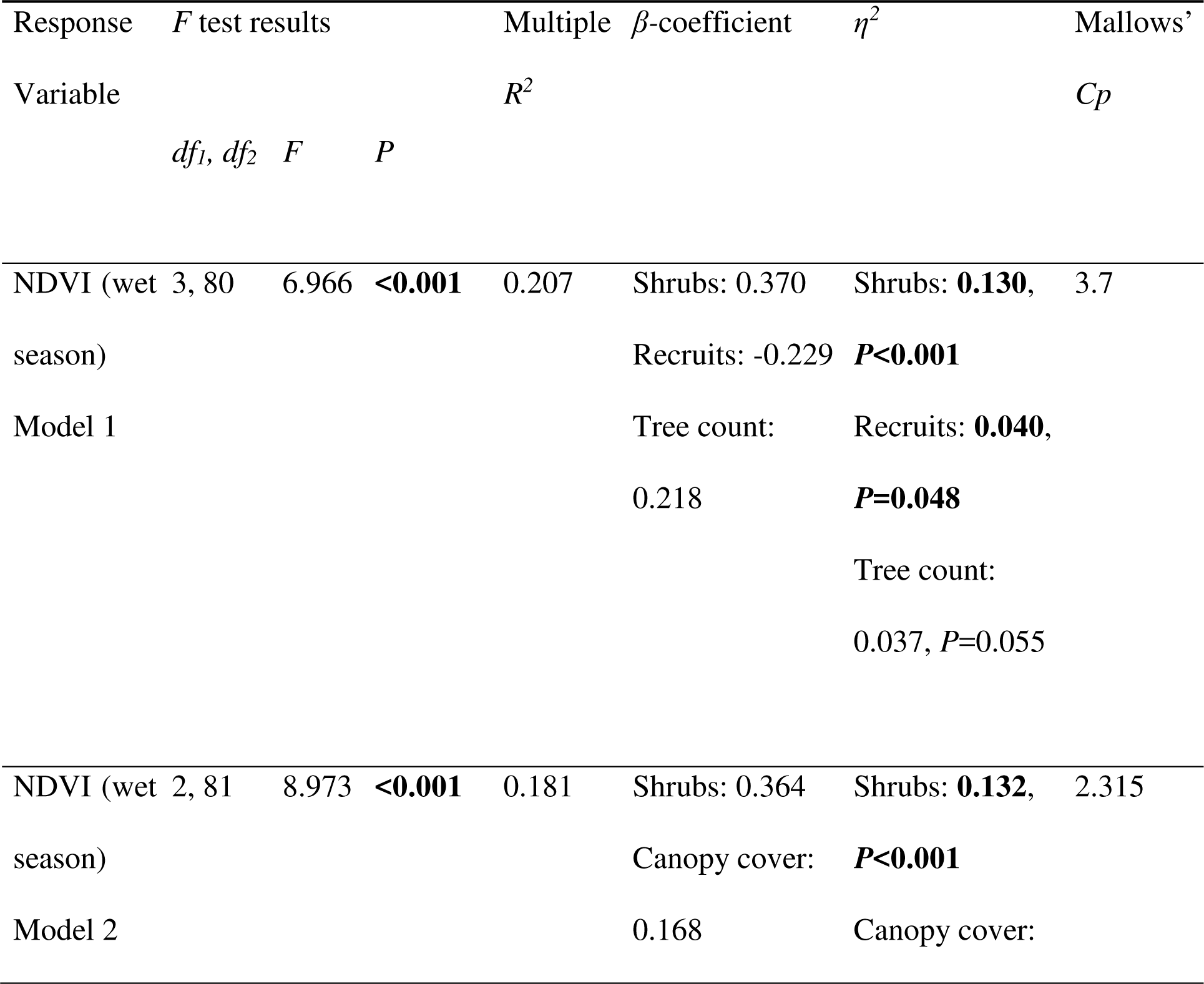
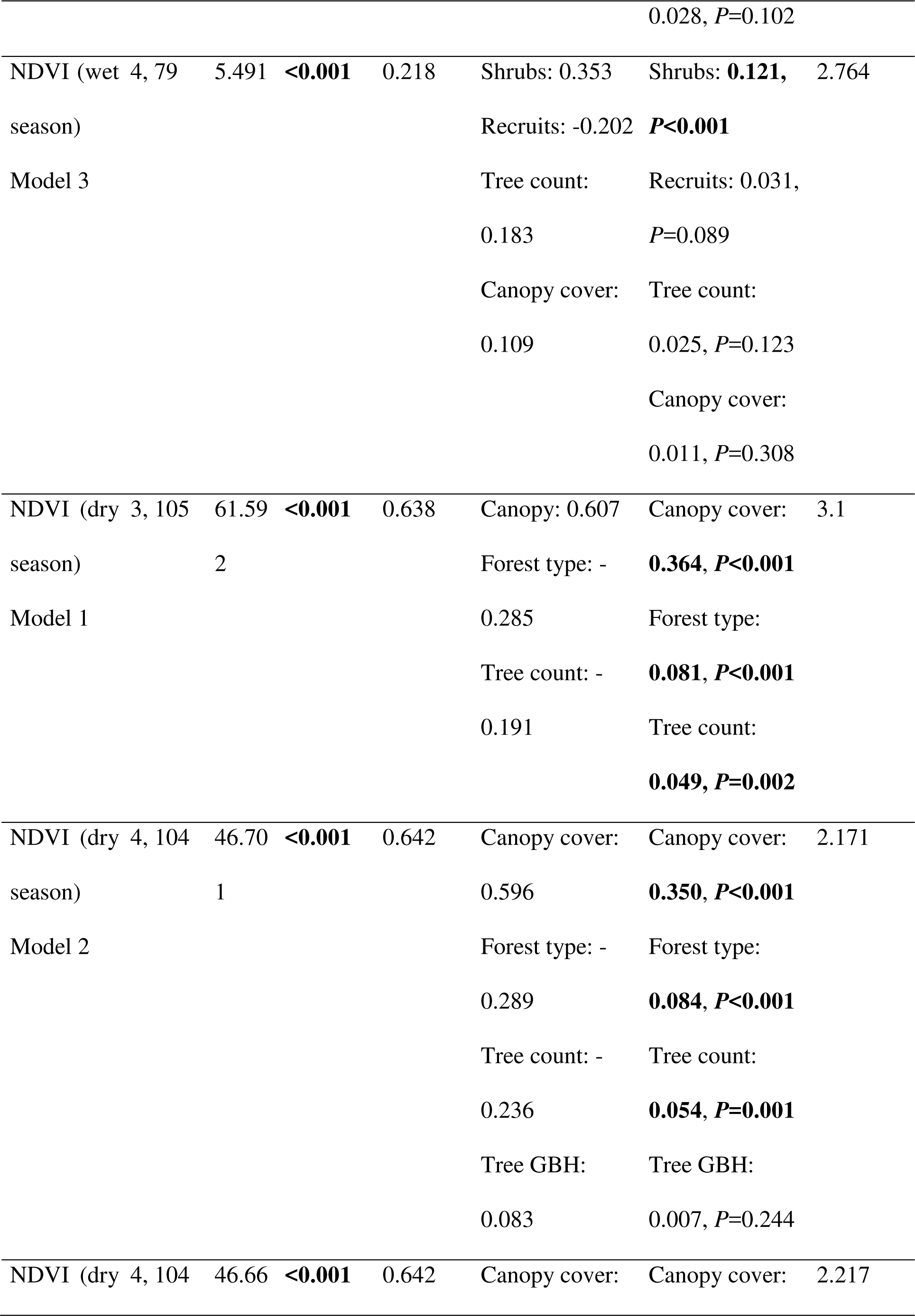
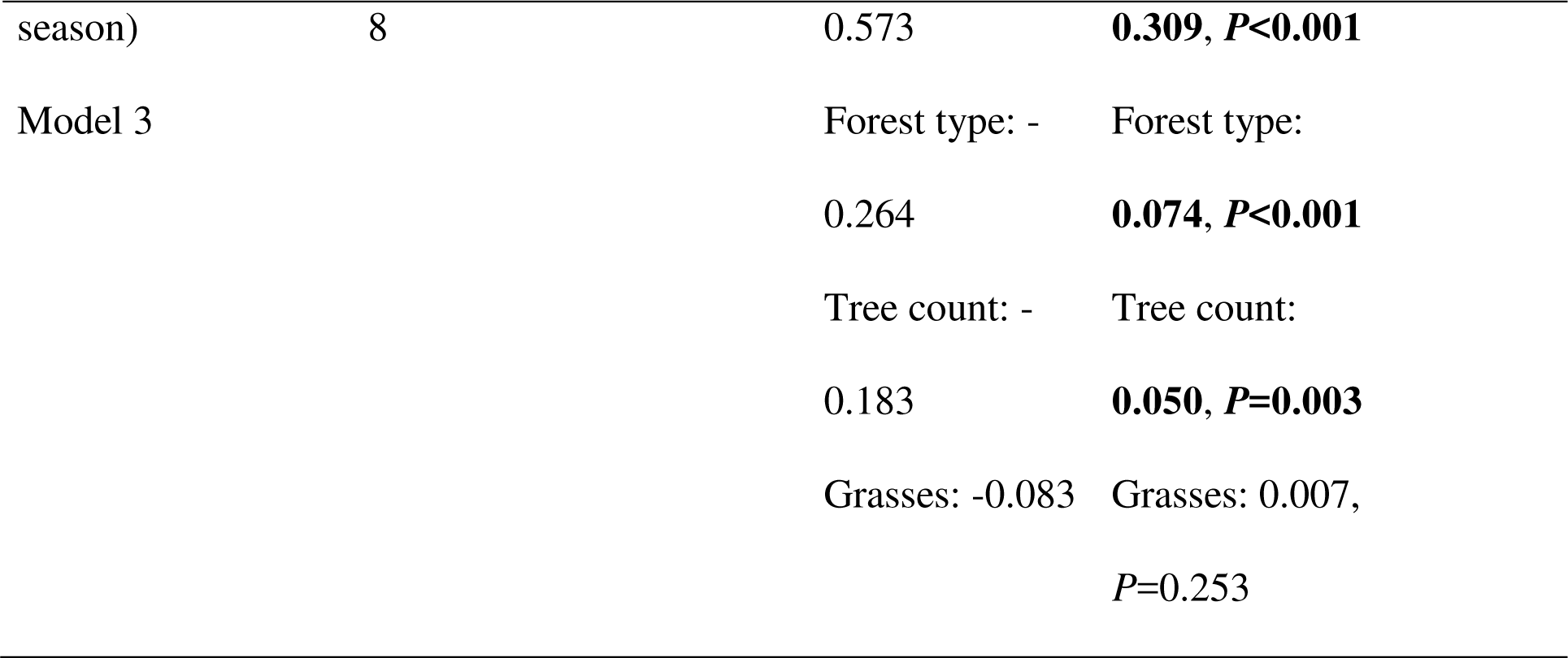
Best-subset regression models of the dependence of NDVI on the abundance of different vegetation categories and canopy cover (all continuous predictors) and forest type (categorical predictor).

The best subset regression model explaining the total abundance of graminoids in the wet season consisted of three variables - forest type, canopy cover and abundance of shrubs, which explained 43.2% of the variation in the total abundance of graminoids. In the dry season, total graminoid abundance was best explained by a model which consisted of six explanatory variables - forest type, canopy cover, abundance of herbs, shrubs, recruits and tree count, which explained 38.2% variation in graminoid abundance. Canopy cover had a negative effect on total graminoid abundance in both the seasons, and its effect relative to those of other vegetation components was larger. The top three models for each season are shown in Table 8.

**Table 8.**
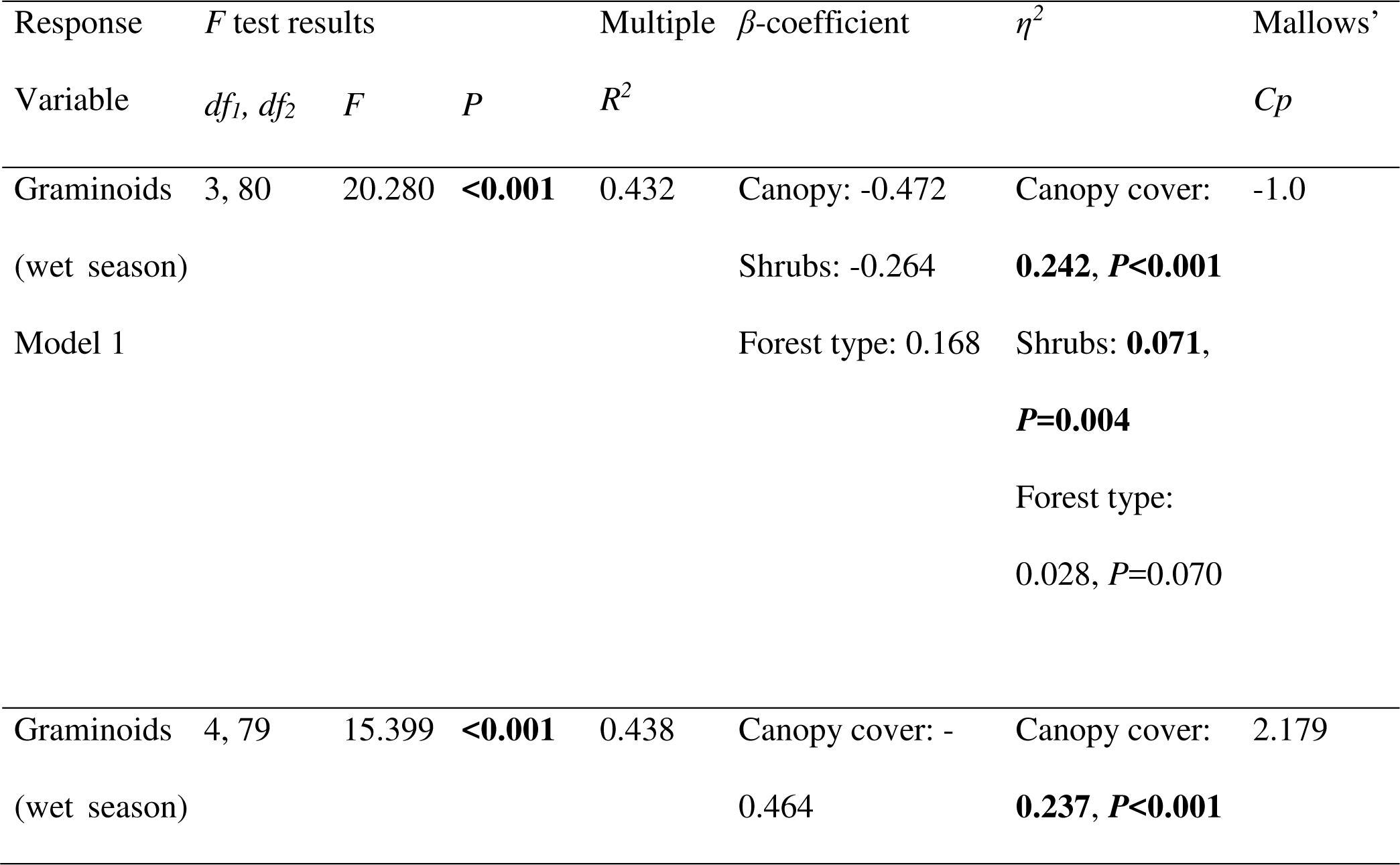
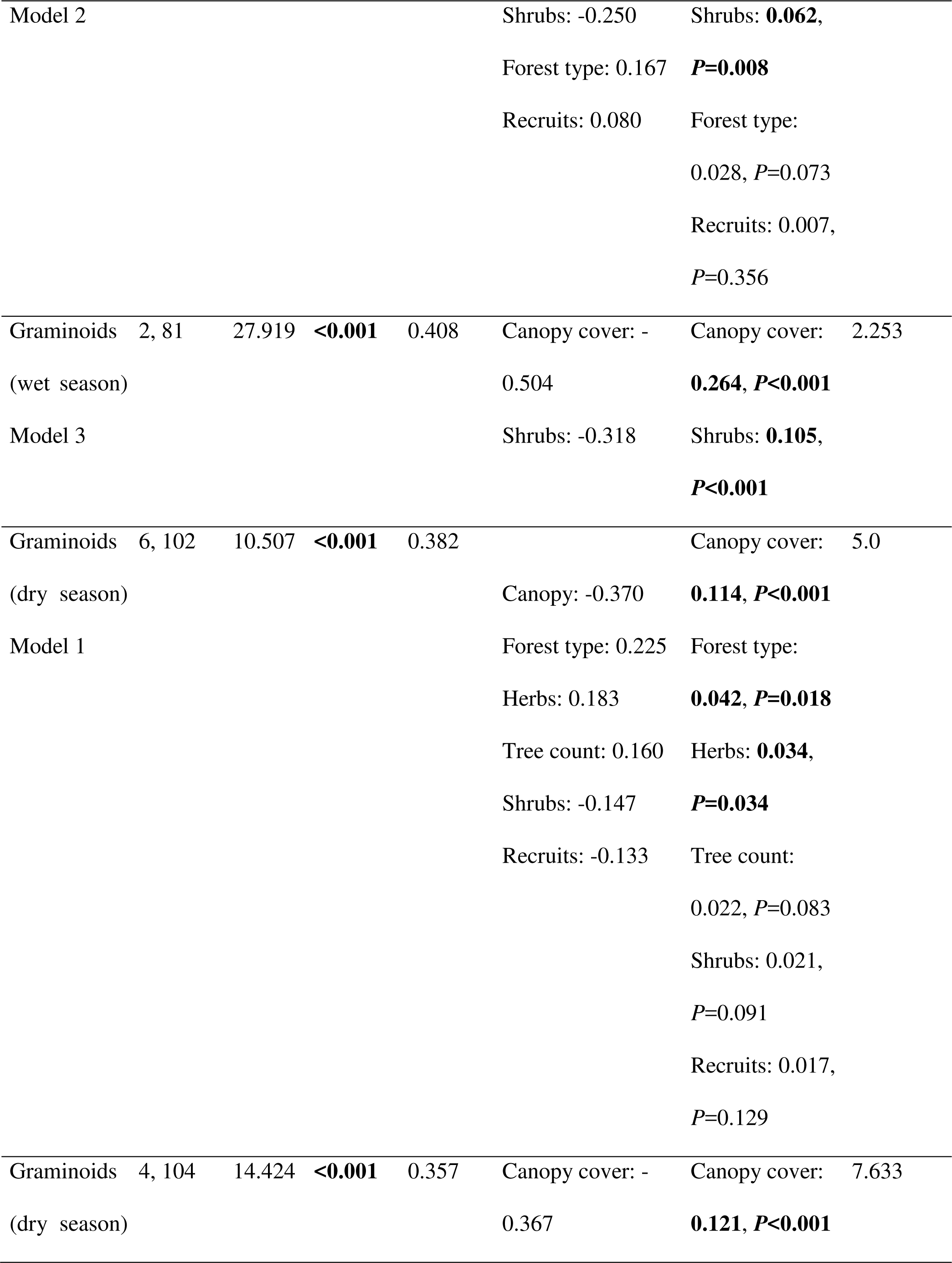
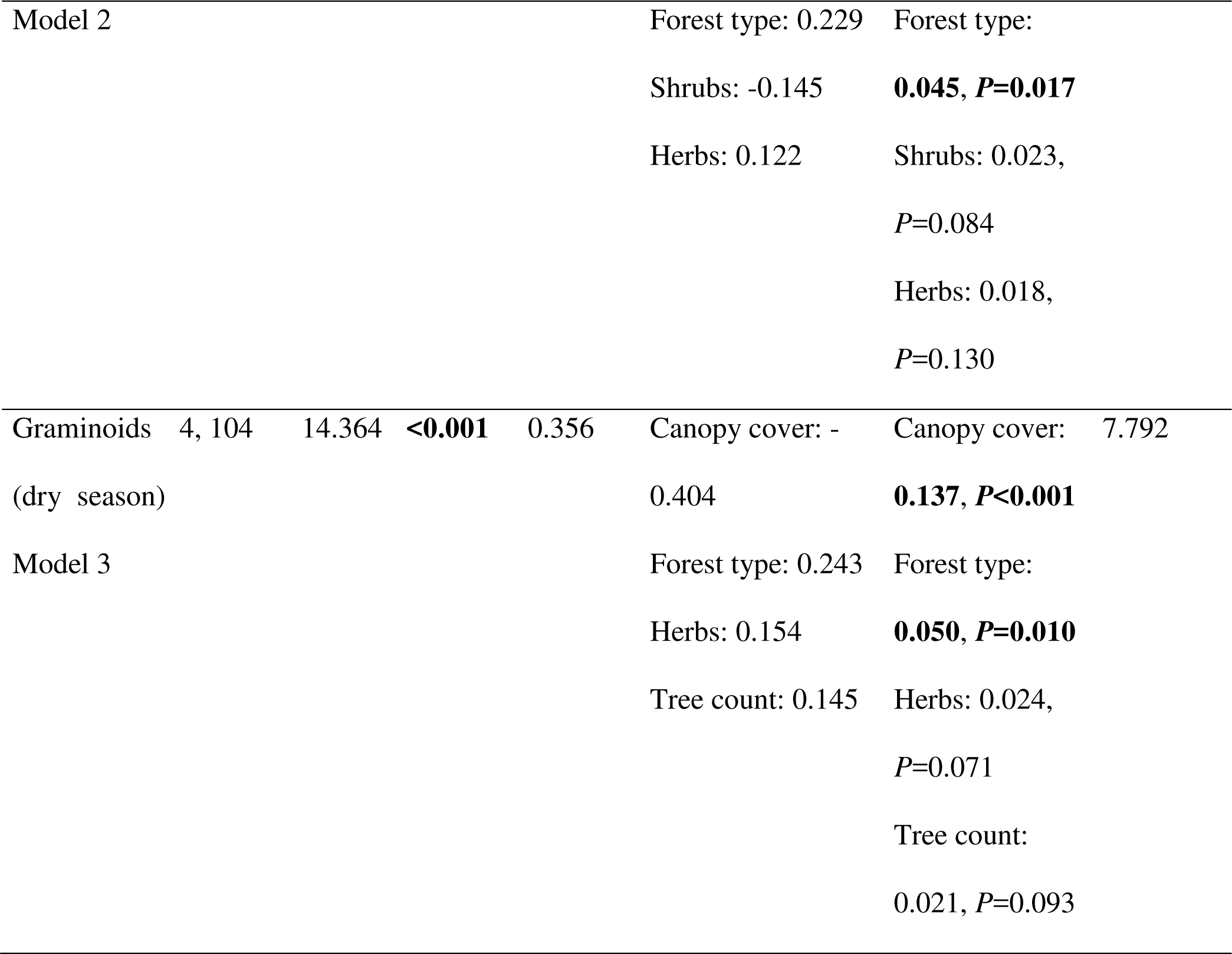
Best subset regression models explaining the dependence of total graminoid abundance on other vegetation variables (continuous predictors) and forest type (categorical predictor).

**3. Comparing NDVI versus spatial interpolation in mapping graminoid abundance**

The raster surfaces obtained from spatial interpolation of field data on graminoids explained significant variation in graminoid abundance. The variation in total graminoid abundance of the 50 dry season verification plots explained by kriging (mean*R*^2^=0.491, SD=0.078)was greater than the variation explained by NDVI(mean *R*^2^=0.331, SD=0.060, negative relationship between NDVI and graminoid abundance) (Wilcoxon matched-pairs test: *T*=0.00, *Z*=2.803, N=10, *P*<0.05; see Figure 2 for bar-graph of *R*^2^ values). Based on the 40 wet season verification plots, while NDVI did not have a significant relationship with total graminoid abundance (mean *R*^2^=0.032, SD=0.022), kriging explained a larger variation (mean *R*^2^=0.174, SD=0.113) in total graminoid abundance (Wilcoxon matched-pairs test: *T*=3.00, *Z*=2.497, N=10, *P*<0.05; Figure 2). Similar analyses on the abundance of food graminoids in the dry season showed that both NDVI (mean *R*^2^=0.279, SD=0.067) and kriging (mean *R*^2^=0.309, SD=0.089) explained significant variation in food graminoid abundance but they were not significantly different from each other (Wilcoxon matched-pairs test: *T*=19.00, *Z*=0.866, N=10, *P*=0.386). Based on wet season data, neither NDVI (mean *R*^2^=0.005, SD=0.005) nor kriging (mean *R*^2^=0.030, SD=0.022) explained significant variation in food graminoid abundance in any of the 10 random verification datasets (Figure 2). Maps of kriging estimates of total graminoid abundance and food graminoid abundance are shown in Figure 4 for one random dataset from the dry season, along with an NDVI map (see Supplementary material Appendix 4 Fig. A3 for the wet season).

**Figure 2.**
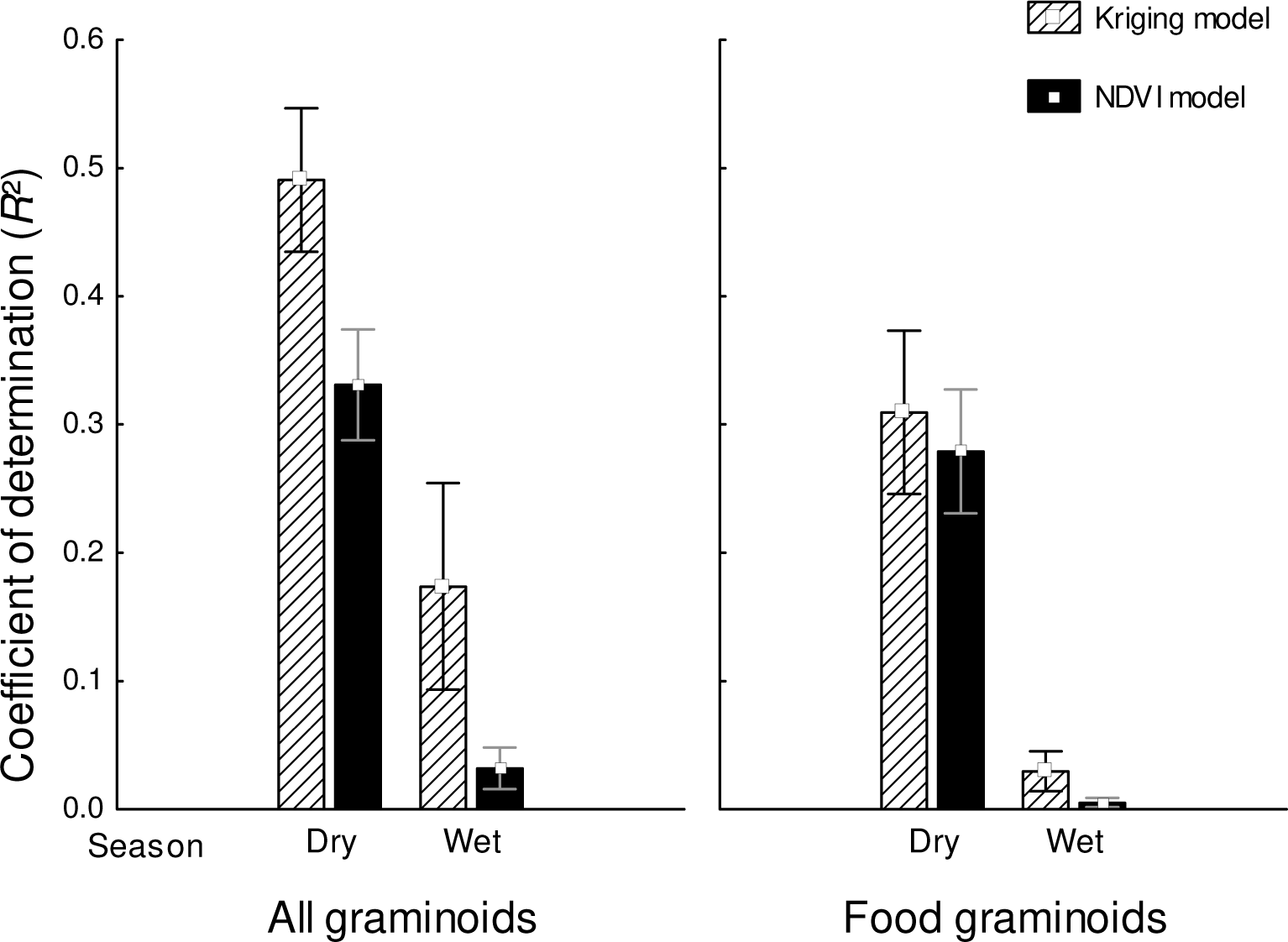
Mean and 1.96 S.E. of coefficient of determination (*R*^2^) obtained from the two types of regression models of graminoid abundance, NDVI and kriging, in the dry and wet seasons. *R*^2^ is shown for both total graminoid abundance as well as elephant food graminoid abundance.

When kriging models obtained from dry season training datasets of two different sample sizes (n=55 and 75 plots) were compared, increasing the sample size from 55 to 75 did not significantly improve the coefficient of determination either for total graminoid abundance (mean *R*^2^=0.491, SD=0.078 for 55 plot sets, mean *R*^2^=0.498, SD=0.074 for 75 plot sets) or for food graminoid abundance (mean *R*^2^=0.309, SD=0.089 for 55 plot sets, mean *R*^2^=0.134, SD=0.070 for 75 plot sets; see Figure 3).

**Figure 3.**
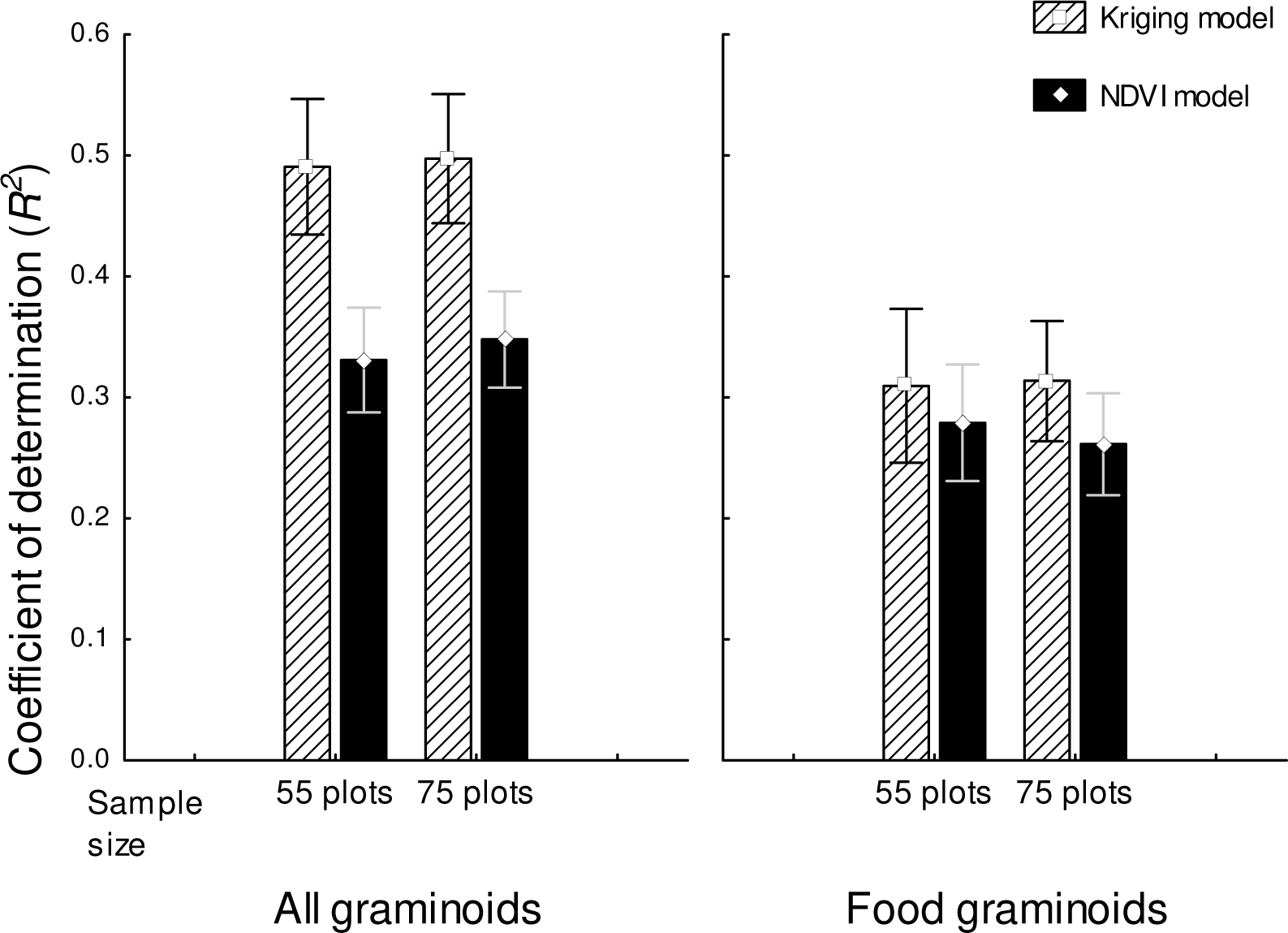
Mean and 1.96 S.E. of coefficient of determination obtained from spatial interpolation (kriging) of total and elephant food graminoid abundance in the dry season using training datasets of 55 plots (50 verification plots) and 75 plots (30 verification plots). Values for NDVI models are also presented for comparison.

**Figure 4.**
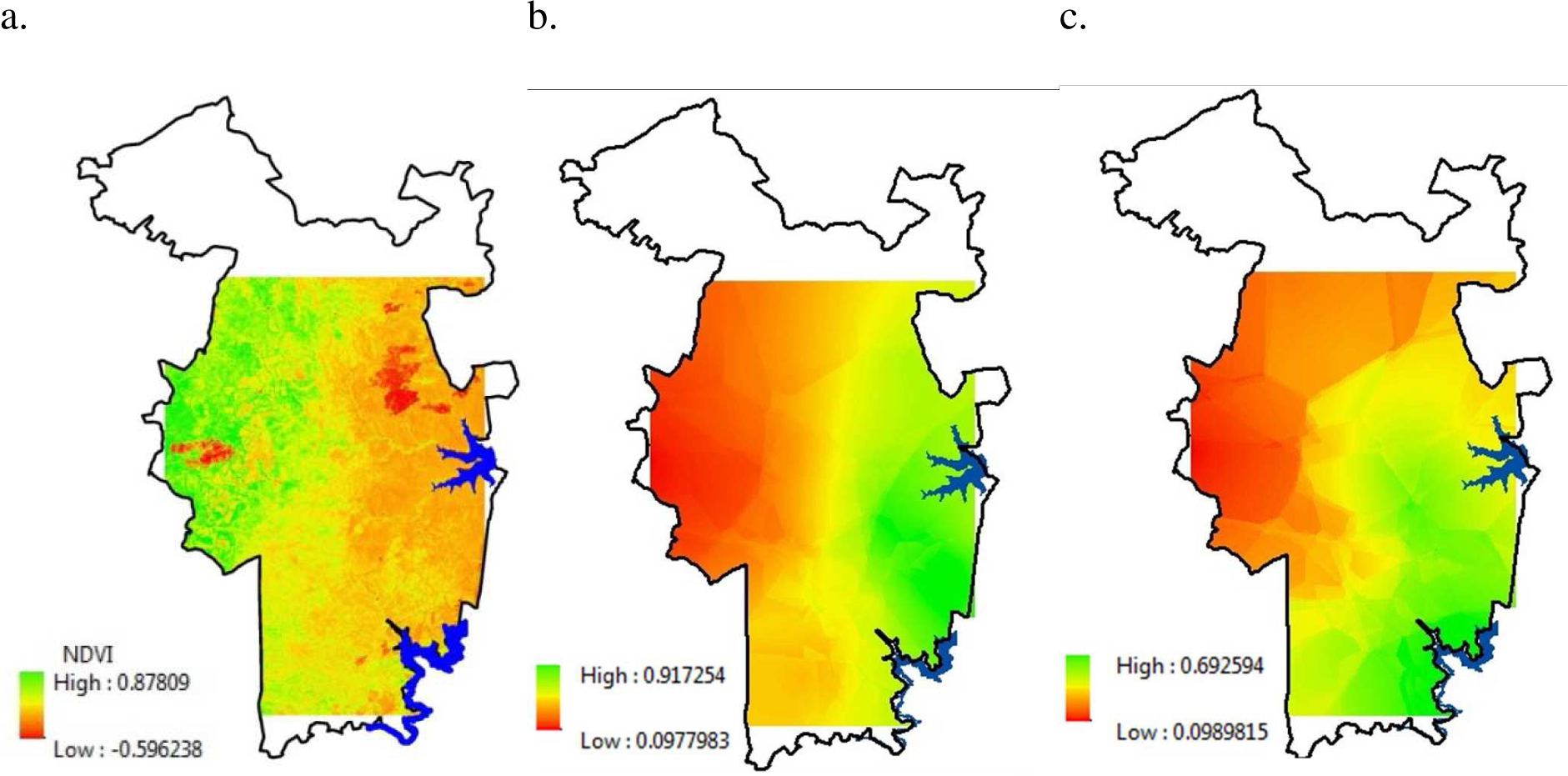
Maps showing (a) NDVI, (b) kriging (spatial interpolation) model of total graminoid abundance, and (c) kriging model of food graminoid abundance in dry season (see Supplementary material Appendix 4 Fig. A3 for wet season). The red patches in the NDVI map are areas affected by forest fire.

## Discussion

This is one of the first studies examining the use of NDVI as a proxy for food abundance of herbivores in tropical forest habitat. We found that NDVI was not a good predictor of elephant forage in any vegetation category in Nagarahole National Park. This ensued both from the small proportion of all plant species that formed elephant food species, as well as from the complexity of multistorey vegetation structure resulting in NDVI poorly capturing total vegetation abundance. The proportion of food species was low overall (13.79%), although the proportional abundance of food species was high within the graminoid category (~80%). The relationship between total abundance and elephant food species abundance was also stronger in the graminoid category than the herb and shrub categories. However, despite the greater proportional abundance of food graminoid species and the ability of total abundance to predict food species abundance to some extent in the graminoid category, NDVI was not a good measure of food graminoid abundance because of the multi-storey vegetation structure in the study area giving rise to a negative relationship between NDVI and total graminoid abundance (see below). NDVI was also not a good measure of food species abundance in the recruit and tree categories despite a moderate proportional abundance (>50%) of food plants and a strong relationship between total abundance and food species abundance because there was no significant relationship between NDVI and total abundance in these strata. The non-significant relationship between NDVI and tree abundance may be the result of the variables that we measured at the ground level, i.e. total tree count and GBH, not representing the primary productivity of trees. Although canopy volume, rather than tree count or GBH, can be measured as the primary productivity of trees, tree canopy is not relevant to foraging by elephants and other terrestrial ungulates. It was not surprising that NDVI was not a good predictor of food abundance in the herb and shrub strata, which had a low (<25%) proportional abundance of food species, as well as a low correlation between total abundance and food species abundance (only total abundance of shrubs in the wet season was significantly correlated with NDVI). The low similarity in distribution of food species abundance and total abundance in herbs and shrubs is of relevance because, although grasses dominate the diet, when grasses are scarce in the forest during the dry season, elephants may feed on fruits and branches of herbs and shrubs in addition to bark from trees.

Previous assessments of NDVI as a measure of food abundance or quality for mammals have been carried out only in relatively open habitats (like steppe grassland, rangeland, savannah, etc.), with the exception of two studies in forest habitats (Willems *et al.* 2009, Borowik *et al.* 2013). Willems *et al.* (2009), in a study of vervet monkey ranging in South Africa, found that while NDVI was strongly correlated with leaf abundance (Pearson’s *r=* 0.923), the abundance of relevant food resources (flowers, fruits and seeds) was not linearly correlated with NDVI. However, a good fit (*R*^2^=0.806) of NDVI with relevant food was obtained using a quadratic polynomial curve. Borowik *et al.* (2013) found the relationship between NDVI and ground vegetation biomass in a temperate forest to be positive in spring but negative in summer when shrub and tree leaves were fully developed. Borowik *et al.* (2013) suggested that canopy closure in the productive season may dominate the signals received by satellite sensors and may also negatively affect ground vegetation. Their latter results are similar to the negative relationship between NDVI and graminoid abundance that we observed in both seasons in our study. Some previous studies in open habitats have also not found very high correlations between NDVI and forage abundance or quality. Zengeya *et al.* (2013) found NDVI obtained from multispectral data to explain low to moderate proportions (*R*^2^=0.01 and 0.48) of forage quality (measured by nitrogen concentration) for cattle and wild ungulates in a rangeland in Zimbabwe. Tsalyuk *et al.* (2017) also found only a moderate proportion of variation in total abundance of different vegetation forms in mixed-savannah habitat in Etosha National Park to be explained by average annual NDVI (grass cover: *R*^2^=0.43, shrub cover: *R*^2^=0.30, tree density: *R*^2^=0.34, and tree biomass: *R*^2^=0.37; graminoids in the dry season from our study: *R*^2^=0.301), although much stronger relationships were found based on multi-year NDVIs. In contrast, while studying forage quality and quantity for grazing livestock in steppe grasslands of Inner Mongolia (China), Kawamura *et al.* (2005) found that NDVI explained 53-75% of the variation in total biomass, 54-74% variation in live biomass, and 48-68% variation in standing crude protein.

In tropical forests, primary productivity is often concentrated in the canopy layers (e.g. Thakur *et al.* 2017, see also Roy and Ravan 1996 for biomass), which are beyond the reach of terrestrial animals. We found that canopy cover contributed significantly to NDVI in the dry season, whereas the effects of other vegetation variables, when significant, were small. The difference in the predictability of NDVI from canopy cover and vegetation between the dry and wet seasons *(Multiple R*^2^ in the wet season: 0.21; dry season: 0.64) suggests a saturation of the relationship between NDVI and vegetation variables when the productivity is high, as highlighted previously also (see Pettorelli *et al.* 2011, Leyequien *et al.* 2013). Canopy cover, although positively related to NDVI, negatively affected the abundance of graminoids, the primary forage of elephants and other ungulates (Baskaran *et al.* 2010, Ahrestani *et al.* 2012). Total graminoid abundance was also affected to a smaller extent by other vegetation categories in the upper layers (see Table 8), possibly through shading or allelopathic effects. Thus, the presence of multistorey vegetation seems to result in the negative relationship between NDVI and graminoid abundance (Figure 5). This negative (and somewhat weak) relationship between NDVI and graminoid abundance is likely to be found across other tropical forests also because of a somewhat continuous canopy layer. Although NDVI was not measured, canopy tree density was negatively related to the abundance of elephant food plants in the understorey in the tropical rainforest of Congo (Blake 2002, pp. 38-40).

**Figure 5.**
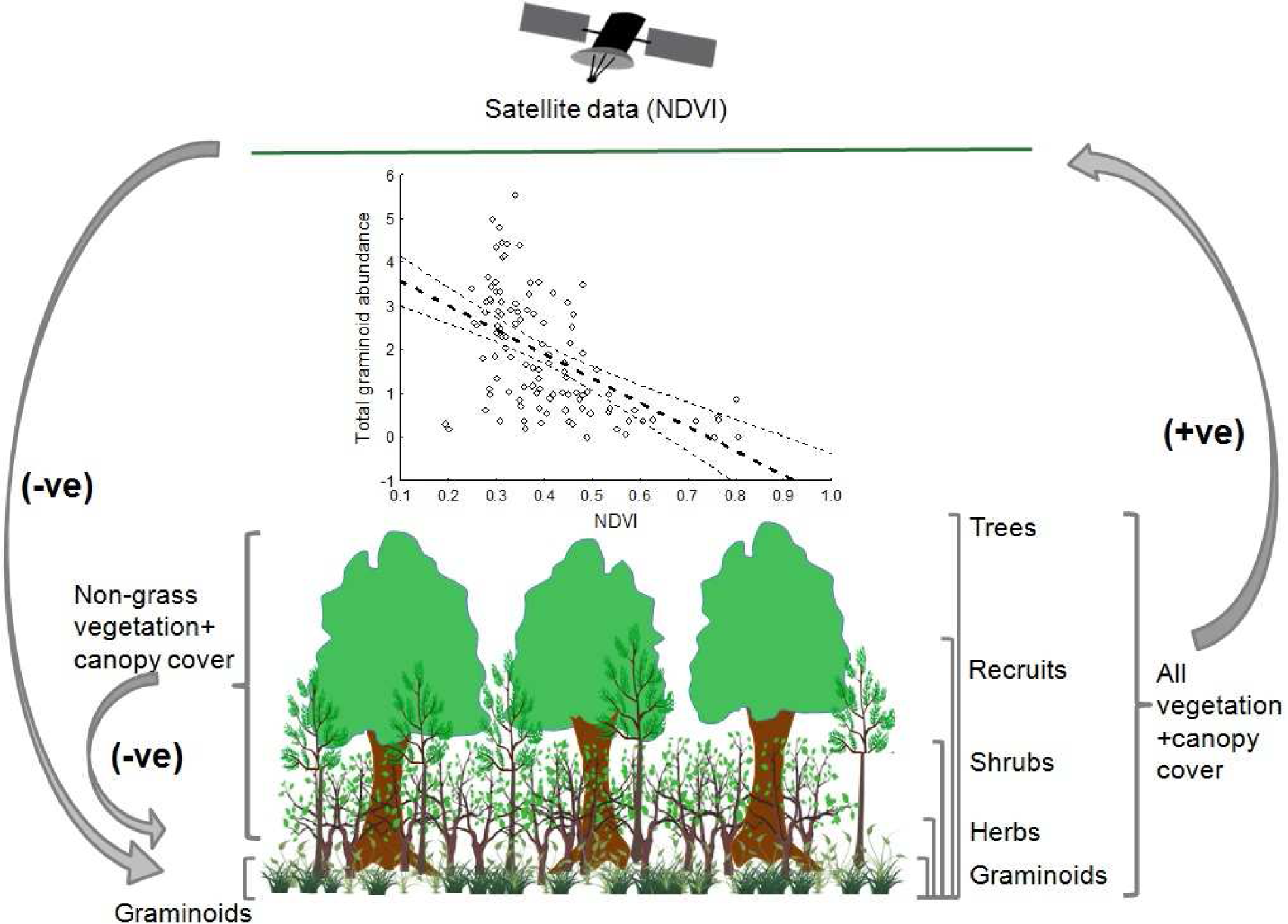
A schematic representation of the relationship between graminoid abundance, non-graminoid vegetation and satellite-derived NDVI productivity.

Based on our results, we caution against the use of NDVI and other remotely-sensed vegetation indices as a proxy for food abundance of herbivores in tropical forests. Although the advantage of rapid mapping of vegetation abundance (Roy and Ravan 1996, Sannier *et al.* 2002, Kawamura *et al.* 2005, Wilson *et al.* 2011) and landscape heterogeneity (e.g. Murwira and Skidmore 2005) has made NDVI a popular habitat covariate in studies of animal ecology and habitat management (see Pettorelli *et al.* 2005, 2011), we find that NDVI is unlikely to be a reliable indicator of food abundance for terrestrial grazing herbivores in tropical forests because of multistorey vegetation interfering with detection of ground vegetation by satellite-based indices, and selectivity of forage by herbivores. While NDVI has been shown to be an informative measure of resource abundance for African elephant populations in savannah habitats (Duffy and Pettorelli 2012), it should not be transferred to a completely different habitat without validation. Studies on Asian elephant habitat use (Rood *et al.* 2010 in Sumatra, Srinivasaiah *et al.* 2012 and Lakshminarayan *et al.* 2015 in southern India, Marasinghe *et al.* 2015 in Sri Lanka) that have used NDVI as a proxy of forage abundance cite papers that either themselves (Pettorelli *et al.* 2011, cited in Lakshminarayan *et al.* 2015) caution against the use of NDVI in densely vegetated areas because the linear relation between NDVI and leaf area index does not hold, or are studies from structurally different vegetation (Hansen *et al.* 2009 based on tundra vegetation, cited in Rood *et al.* 2010; Wittemyer *et al.* 2007, Young *et al.* 2009, based on African savannahs, cited in Srinivasaiah *et al.* 2012) that moreover do not actually examine the relationship between NDVI and animal forage abundance. Moreover, associations between NDVI and animal habitat use may result from correlated responses to other environmental variables such as water availability, protection status/human disturbance, or shade rather than forage abundance, necessitating the teasing apart of these confounding variables from forage abundance. For example, in large, deforested landscapes, preference of areas with high productivity based on NDVI could be interpreted either as a preference for riparian forest habitat or for areas with greater forest cover. Similarly, preference for areas with high NDVI in fragmented habitats could be interpreted as a preference for forage or for safer habitats. Moreover, simultaneous use of multiple remotely-sensed indices that are known to be correlated in deciduous forests (see Madugundu *et al.* 2008) and arbitrary justification regarding what they represent (for example, leaf area index as a proxy of shade and NDVI as a proxy for food in Srinivasaiah *et al.* 2012) makes it difficult to tease apart various effects such as food, human disturbance, and shade on habitat use. NDVI has been used as a proxy for resource availability or habitat quality in forest habitats in the case of other mammalian taxa such as primates *(Papio hamadryas* and *Papio h. anubis* from north-eastern Africa, Zinner *et al.* 2001), ungulates *(Axis kuhlii* and *Muntiacus muntjak* from Indonesia, Rahman *et al.* 2017), and arboreal marsupials *(Petauroides volans, Petaurus breviceps, Pseudocheirus peregrinus, Trichosurus vulpecula, T. cunninghami* from eastern Australia, Youngentob *et al.* 2015) also, and field-based sampling of relevant vegetation in these habitats can validate how useful NDVI is in mapping of forage abundance.

We found significant effects of season, forest type, and transects within forest types, and their interactions on food abundance in different vegetation categories, showing considerable spatial and seasonal heterogeneity in food species abundance for Asian elephants in our study area. Such variability in food abundance is expected to affect the spatio-temporal patterns of movement, foraging, and habitat use, as seen in African elephants (e.g. Blake 2002, Marshal *et al.* 2010). Total abundance or NDVI that do not reflect food species abundance patterns would, therefore, yield misleading results. We suggest that ecologists studying large mammal foraging in tropical forests, directly measure food abundance from field sampling rather than remote sensing unless verification is carried out for the study area or alternate rapid methods are developed. Spatial interpolation (kriging) could explain larger variation in total graminoid abundance than NDVI in our study, although the *R*^2^ remained modest even upon increasing the sample size of training sets (55 to 75 plots). Neither method was very useful in estimating food graminoid abundance, especially during the wet season. Our study area was spread over a few hundred sq. km. and our data were from fine-scale plots, relevant to the spatial scale at which elephants feed. A general regression analyses at the resolution of 1km (rather than 20m × 5 m plots) suggested that our results hold at this coarser resolution (negative effect of NDVI on total graminoids abundance in dry season: *F*[2,19]=9.343, *Multiple R*^2^=0.496, *P*=0.001; *β*_*NDVI*_=-0.542, *P*=0.022; effect not significant in the wet season: *F*[2,14]=1.514, *Multiple R*^2^=0.178, *P*=0.254; *β*_*NDVI*_*=-0.384, P*=0.136). However, at even coarser resolutions in large landscapes with a variety of land cover types, NDVI may be a proxy of habitat suitability, because high NDVI would then be associated with structural aspects of the landscape, for which the focal species may have a preference.

To conclude, our results show that the abundance of non-food vegetation undermines the utility of NDVI as an indicator of elephant food in forest habitats. Canopy in upper strata negatively affects the abundance of graminoids, which are the primary food type for elephants. Since deciduous forests account for more than 65% of all forests in India (Reddy *et al.* 2015) and one-sixth of all forests in South-east Asia (Wohlfart *et al.* 2014), which harbour many endemic and threatened populations of grazing ungulates and other large mammals that largely feed on understorey vegetation and are likely to be more selective feeders than elephants, our results have wider implications for ecological studies in these habitats. These results call for exploration of other methods of rapid assessment of food resources that may reduce field effort in tropical forest habitats.

## Acknowledgements

Ranga, Suresh, Suresh, Shankar, Krishna, Salim and Altaf provided field assistance. We thank Dr. Ravikumar from Foundation for Revitalization of Local Health Traditions (FRLHT), Bangalore, who identified trees, shrubs and herbs, and Dr. Girish Potdar from Yeshwantrao Chavan College of Science, Karad, Maharashtra, who identified graminoid species. We thank the office of the PCCF, Karnataka Forest Department, and the office of the Conservator of Forests and Director, Rajiv Gandhi National Park, Hunsur, for field permits, and various officials and staff of Nagarahole National Park for support at the field site.

## Statement of authorship

HG and TNCV designed the methodology, HG, EA, and MK collected field data, HG did the analyses with major inputs from TNCV, HG and TNCV wrote the manuscript and all authors read and approved the full contents of the final draft.

## Supplementary material

### Appendix 1

**Table A1:**
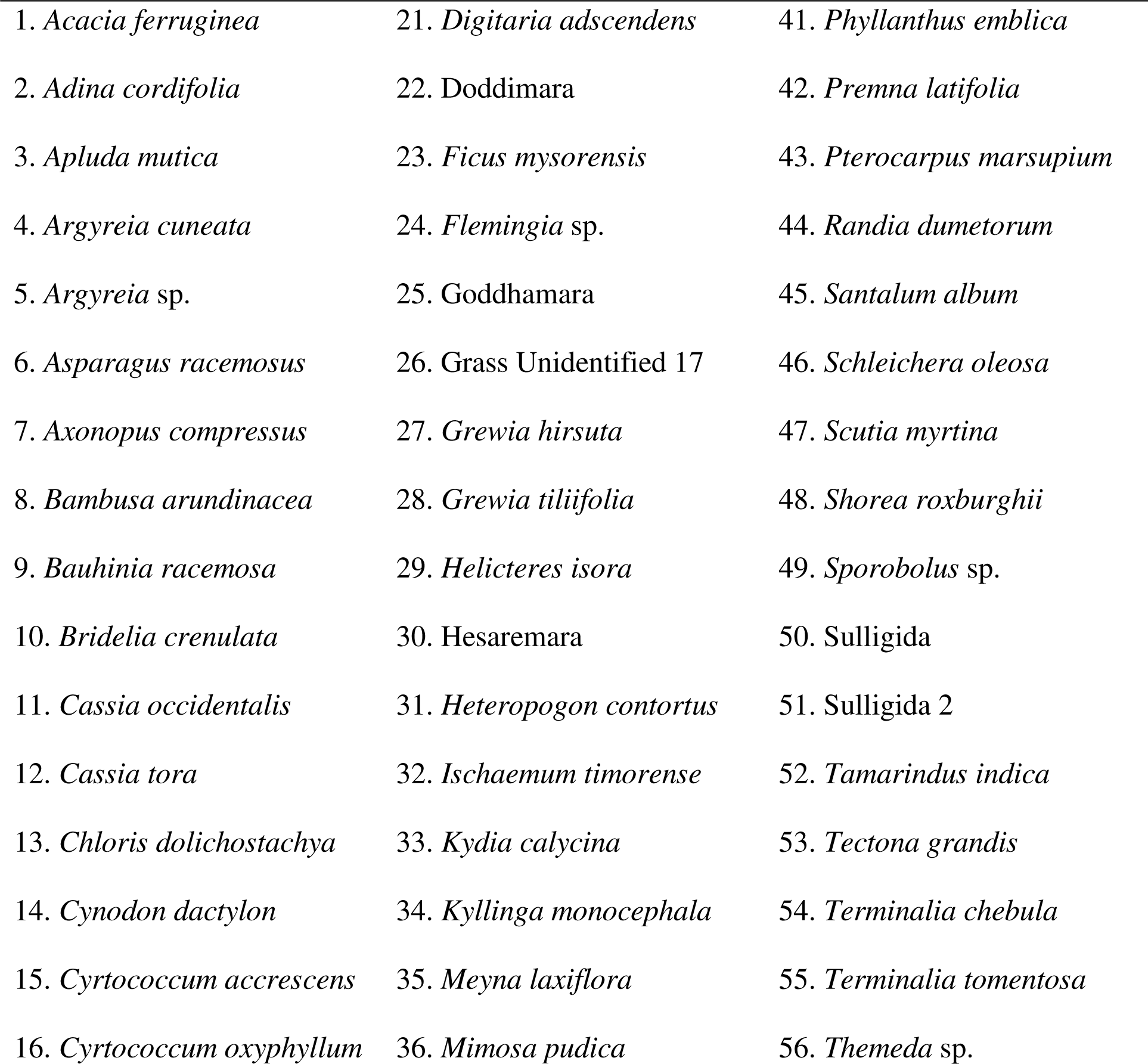
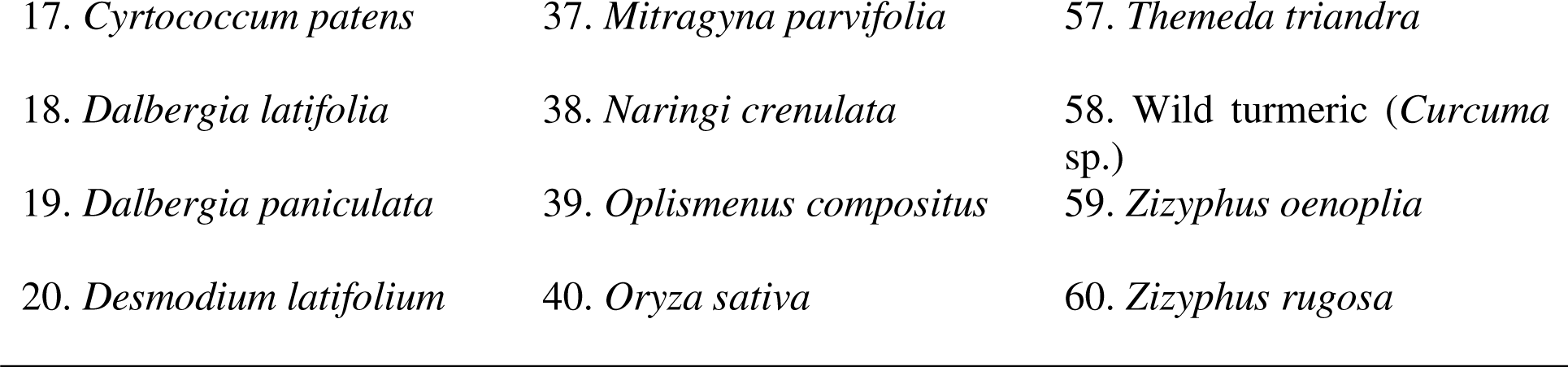
List of elephant food species found in the vegetation plots. Some food species in the list are based on earlier studies from Mudumalai Tiger Reserve (Sivaganesan 1991, Baskaran et al. 2010) and Wayanad Wildlife Sanctuary (Easa 1999), both in the Nilgiris- Eastern Ghats landscape, and close to Nagarahole National Park. Plants that could not be identified are referred to by their local name in most cases.

### Appendix 2

**Table A2:**
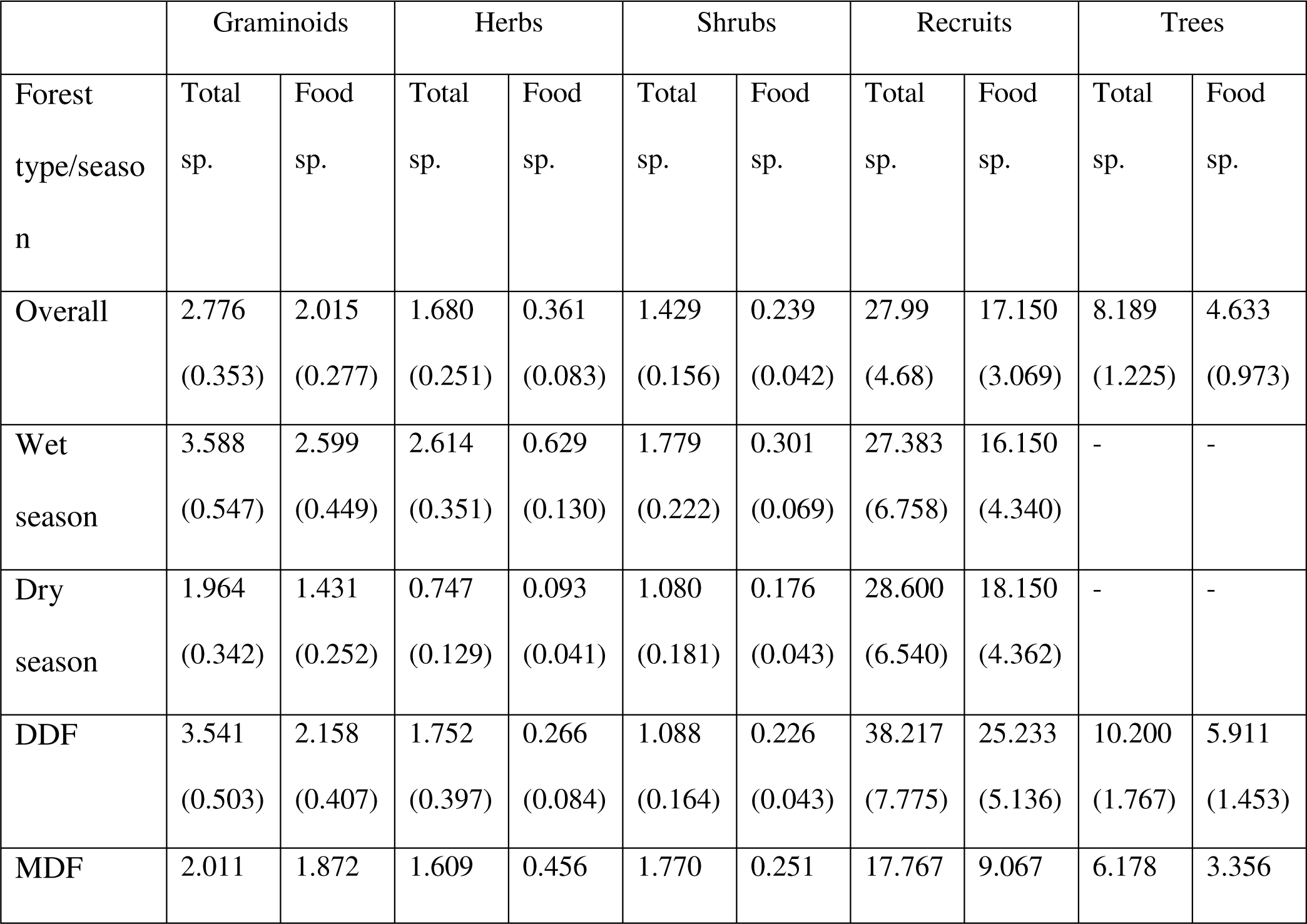
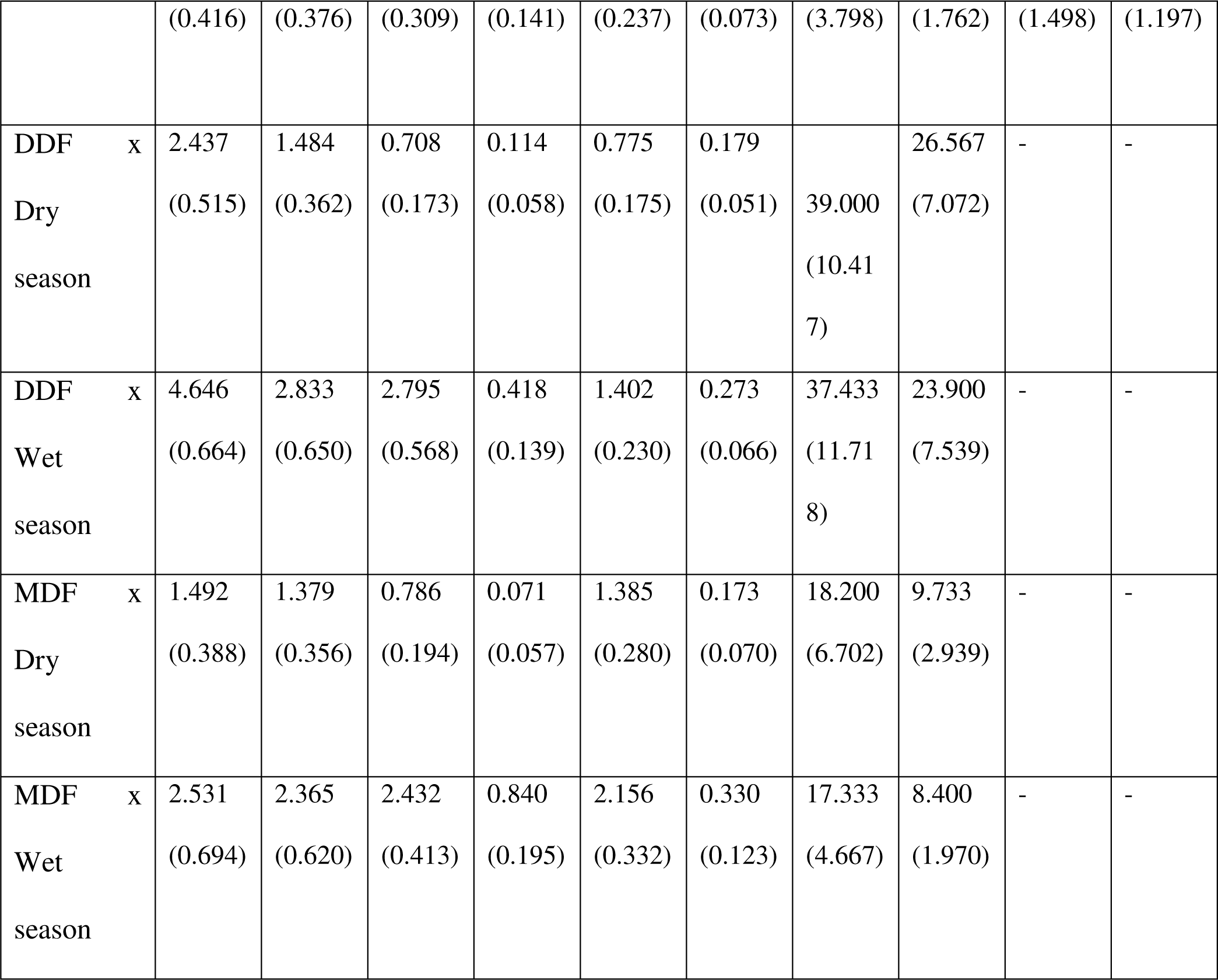
Mean and 1.96 S.E. (within parentheses) total (sum of all species) abundance and food species (sum of food species) abundance in respective vegetation strata for the ANOVA tests shown in Table 3. Values are calculated for different seasons and forest types, and their combinations.

### Appendix 3

**Figure A1:**
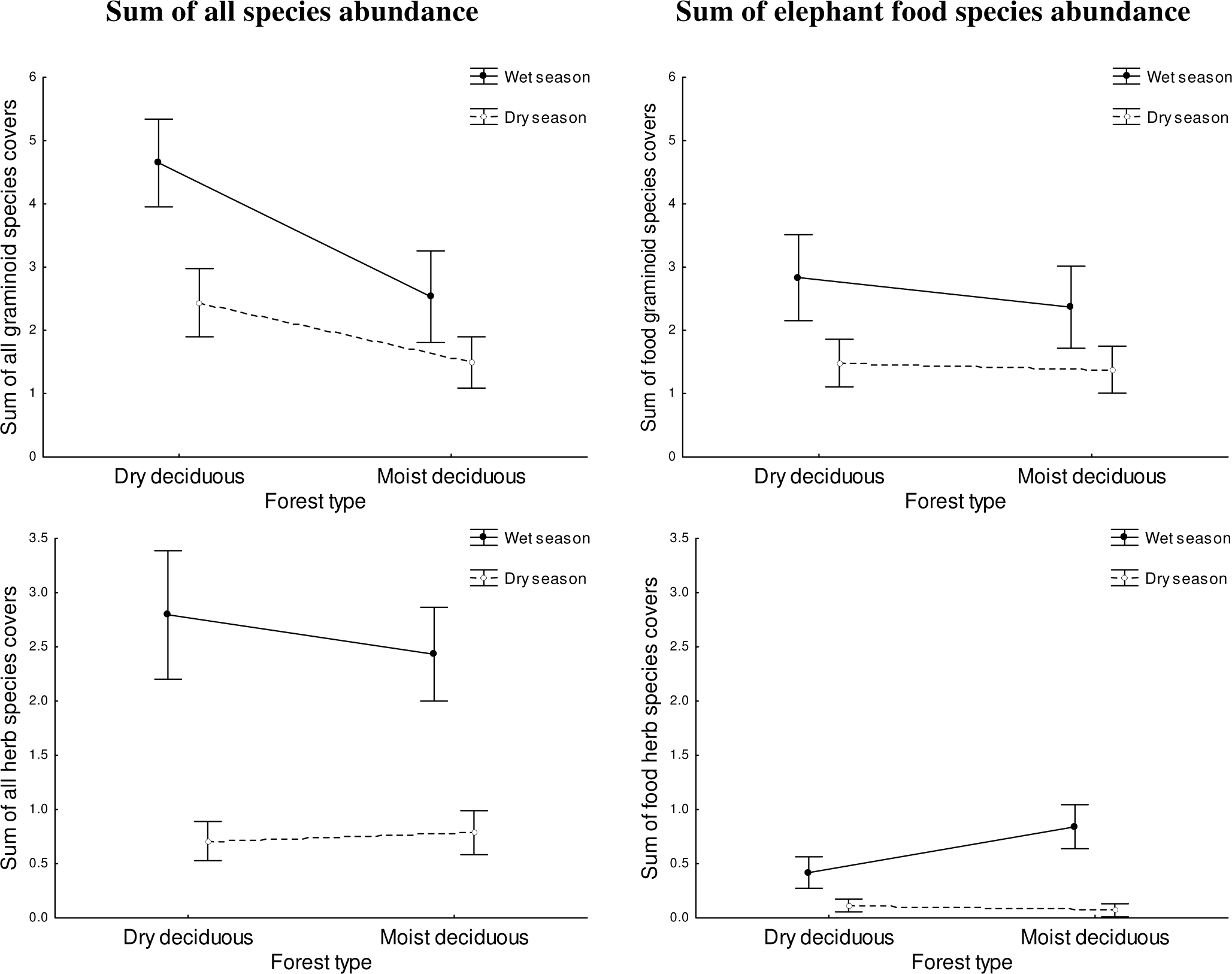
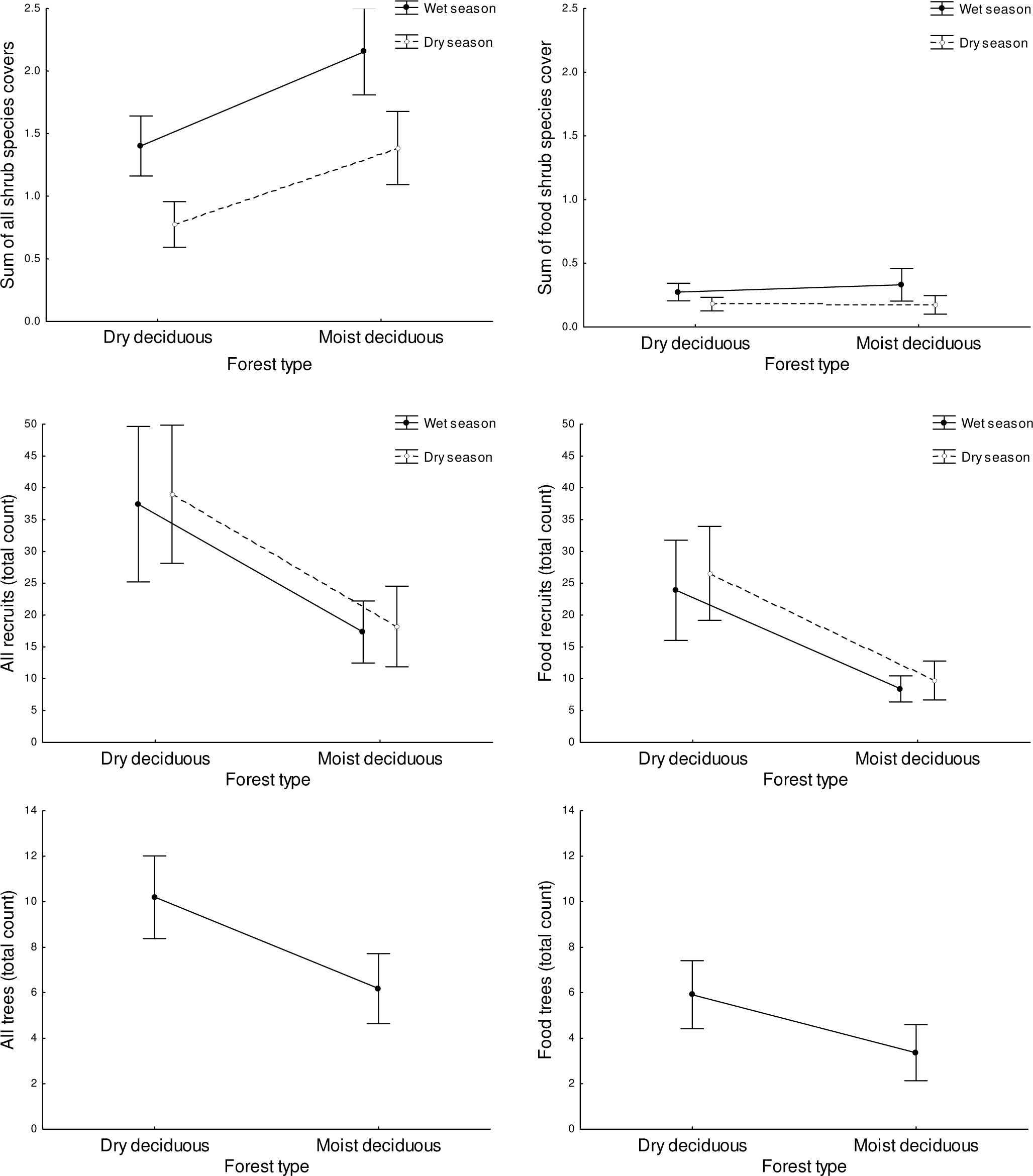
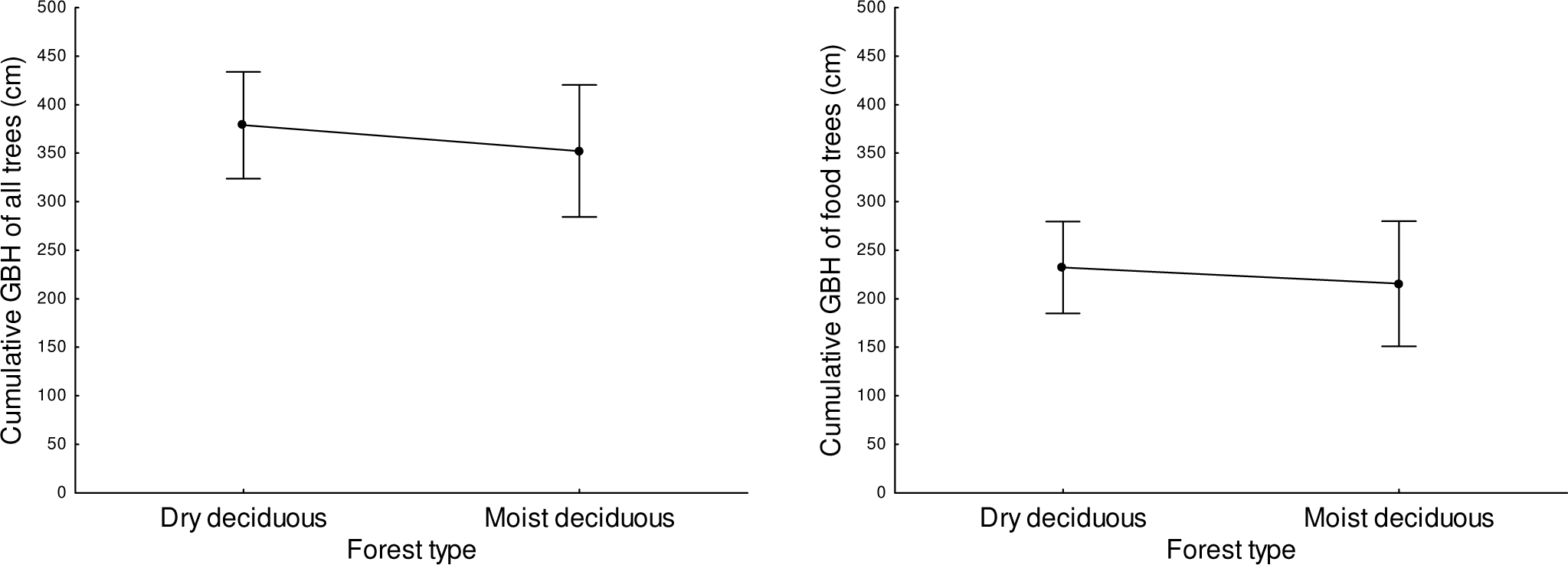
Graph showing mean and 95% CI total and food species abundance in respective vegetation strata for the ANOVA tests shown in Table 3. Values are calculated for pooled data and for different seasons and forest types, and their combinations.

**Figure A2:**
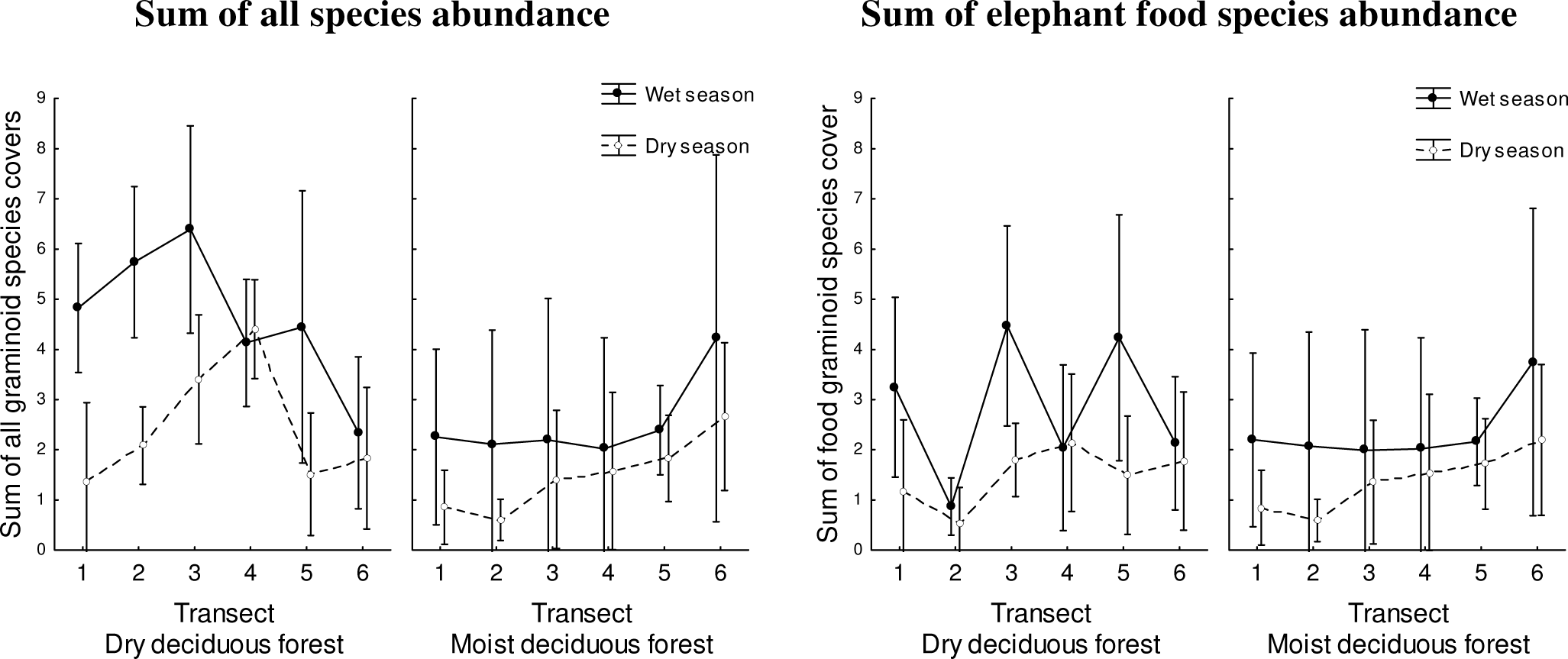
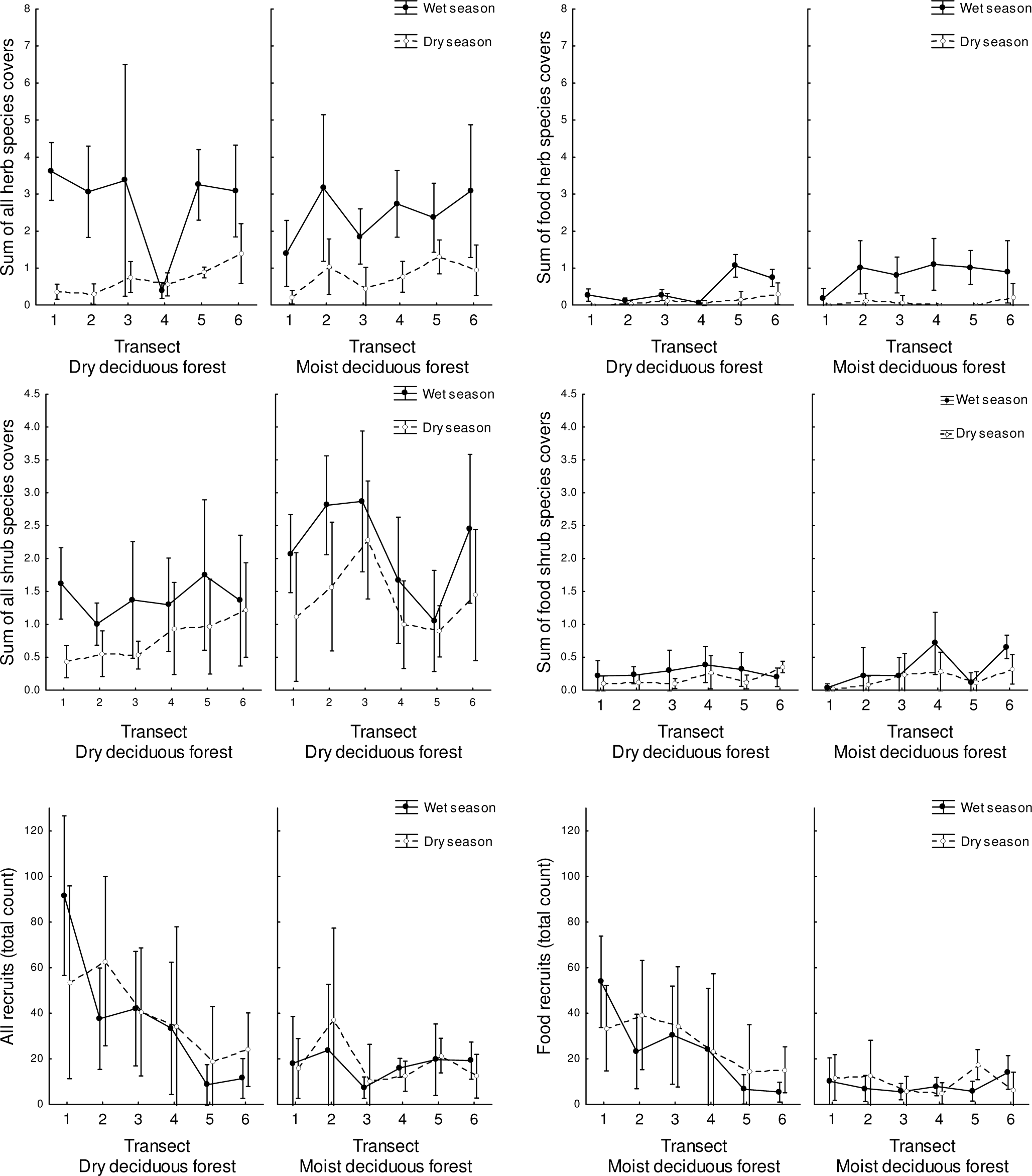
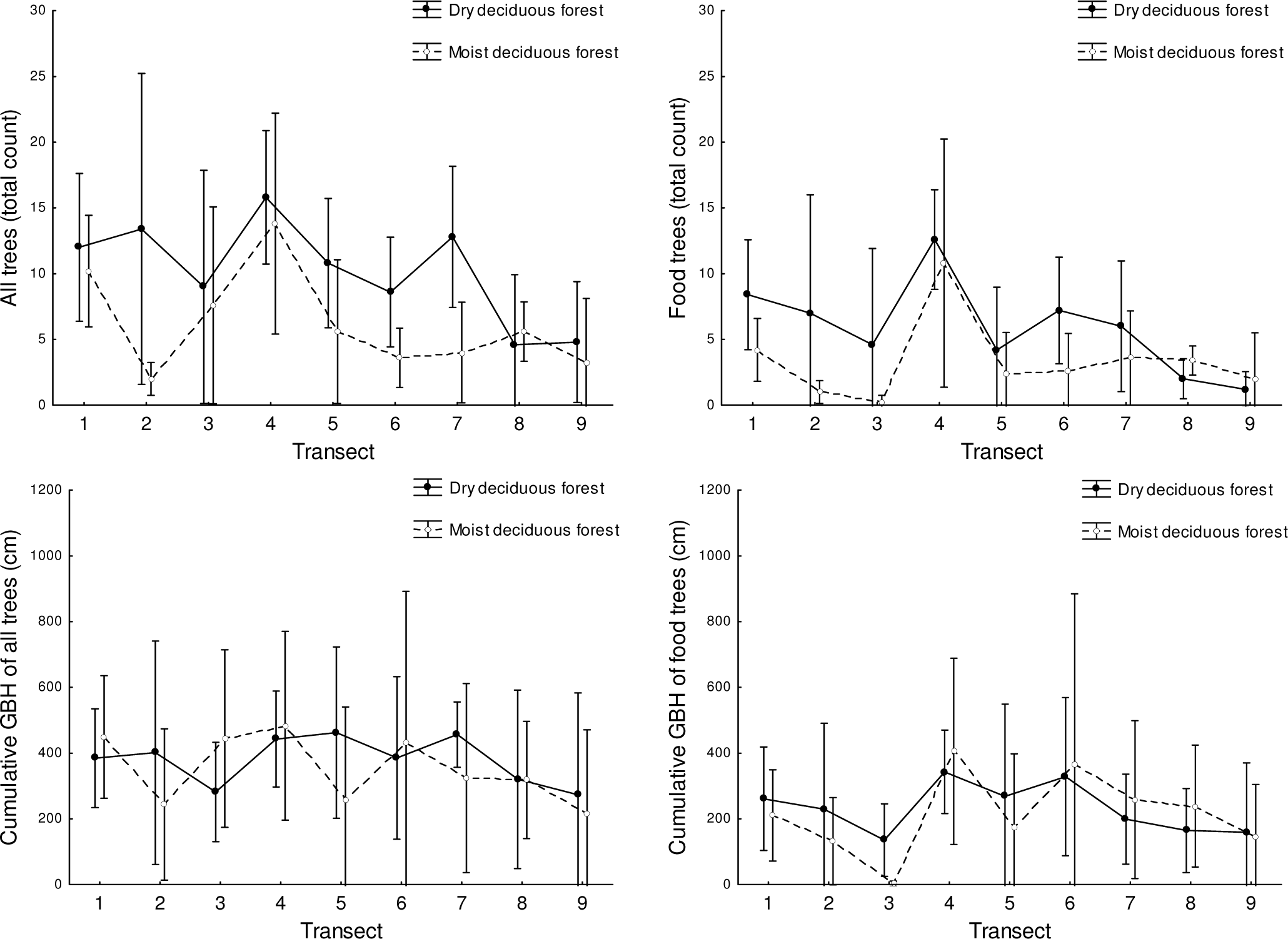
Plot of sum of food species abundance and sum of all species abundance in different transects nested in dry and moist deciduous forests in the two seasons.

### Appendix 4

**Figure A3:**
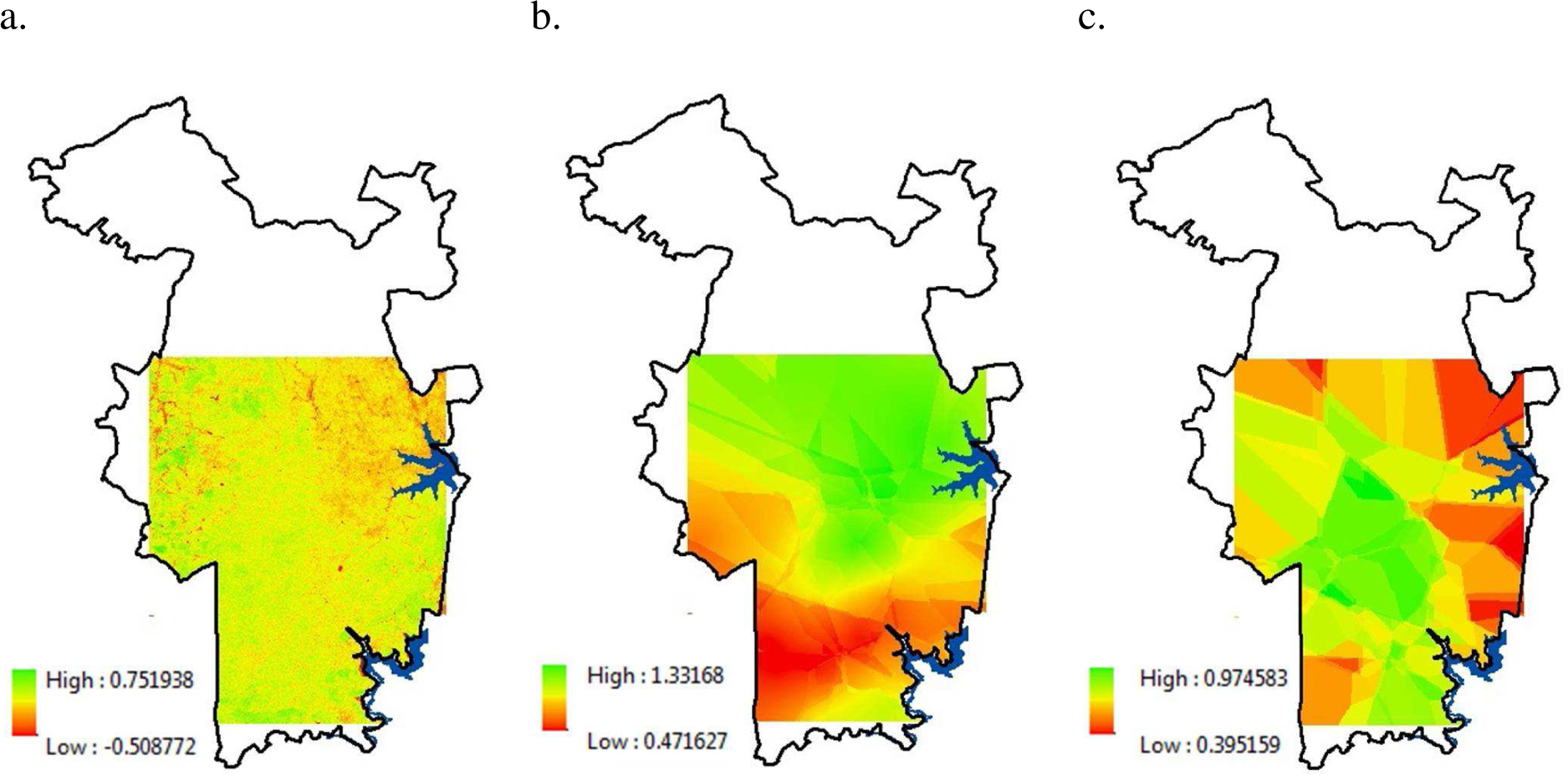
Maps showing (a) NDVI, and kriging (spatial interpolation) models of (b) total graminoid abundance and (c) food graminoid abundance in the wet season.

## References

1. Ahrestani, F.S. et al. 2012. Diet and habitat-niche relationships within an assemblage of large herbivores in a seasonal tropical forest. - J. Trop. Ecol. 28: 385–394.

2. Asian Elephant Research and Conservation Centre. 2006. Southern India Elephant Census 2005. Asian Elephant Research and Conservation Centre, Bangalore, India.

3. Baskaran, N. et al. 2010. Feeding ecology of the Asian Elephant Elephas maximus in the Nilgiri Biosphere Reserve, Southern India. - J. Bomb. Nat. Hist. Soc. 107: 3–13.

4. Belovsky, G.E., 1978. Diet optimization in a generalist herbivore: the moose. - Theor. Popul. Biol. 14: 105–134.

5. Blake, S. 2003. The ecology of forest elephant distribution and its implications for conservation. Doctoral dissertation. - Univ. of Edinburgh.

6. Bohrer, G. et al. 2014. Elephant movement closely tracks precipitation-driven vegetation dynamics in a Kenyan forest-savanna landscape. - Mov. Ecol. 2: 1–12.

7. Borowik, T. et al. 2013. Normalized difference vegetation index (NDVI) as a predictor of forage availability for ungulates in forest and field habitats. - Eur. J. Wildlife Res. 59: 675–682.

8. Das, S. and Singh, T.P. 2016. Forest type, diversity and iomass estimation in tropical forests of estern Ghat of Maharashtra using geospatial techniques. - Small-scale For. 15: 517–532.

9. Demment, M.W. 1982. The scaling of ruminoreticulum size with body weight in East African ungulates. - Afr. J. Ecol. 20: 43–47.

10. Doncaster, C.P. and Davey, A.J. 2007. Analysis of variance and covariance: how to choose and construct models for the life sciences. Cambridge Univ. Press.

11. du Toit, J.T. and Yetman, C.A. 2005. Effects of body size on the diurnal activity budgets of African browsing ruminants. - Oecologia 143: 317–325.

12. Duffy, J.P. and Pettorelli, N. 2012. Exploring the relationship between NDVI and African elephant population density in protected areas. - Afr. J. Ecol. 50: 455–463.

13. E. S. R. I. 2012. ArcGIS desktop: release 10.1. Environmental Systems Research Institute, Redlands, California.

14. Easa, P. S. 1999. Status, habitat utilization and movement pattern of large mammals in Wayanad Wildlife Sanctuary. -Research Report 173, Kerala Forest Research Institute, Peechi, India.

15. Fritz, C.O. et al. 2012. Effect size estimates: current use, calculations, and interpretation. - J. Exp. Psychol. Gen. 141: 2–18.

16. Gautam, H. et al. 2017. Using visual estimation of cover for rapid assessment of graminoid abundance in forest and grassland habitats in studies of animal foraging. - Phytocoenologia 47: 315–327.

17. Hamel, S. et al. 2009. Spring normalized difference vegetation index (NDVI) predicts annual variation in timing of peak faecal crude protein in mountain ungulates. - J. Appl. Ecol. 46: 582–589.

18. Hanley, T.A. 1982. The nutritional basis for food selection by ungulates. - J. Range. Manage. 35: 146–151.

19. Hansen, B. B. 2009. Functional response in habitat selection and the tradeoffs between foraging niche components in a large herbivore. - Oikos 118: 859–872.

20. He, K.S. et al. 2015. Will remote sensing shape the next generation of species distribution models? - Remote Sens. Ecol. Conserv. 1: 4–18.

21. Hedges, S. et al. 2008. Range wide mapping workshop for Asian elephants (Elephas maximus) Cambodia. October 2008. - Report to the U.S. Fish & Wildlife Service.

22. Johnson, C.J. et al. 2001. Foraging across a variable landscape: behavioral decisions made by woodland caribou at multiple spatial scales. - Oecologia 127: 590–602.

23. Kawamura, K. et al. 2005. Comparing MODIS vegetation indices with AVHRR NDVI for monitoring the forage quantity and quality in Inner Mongolia grassland, China. - Grassl. Sci. 51: 33–40.

24. Kazmin, V.D. et al. 2011. Current state of forage resources and feeding of reindeer (Rangifer tarandus) and musk oxen (Ovibos moschatus) in the arctic tundras of Wrangel Island. - Biol. Bull. 38: 747–753.

25. Lakshminarayanan, N. et al. 2015. Determinants of dry season habitat use by Asian elephants in the Western Ghats of India. - J. Zool. 298: 169–177.

26. Leica Goesystems. 2005. ERDAS Imagine 9.1. Leica Geosystems Geospatial Imaging, Norcross, GA.

27. Leyequien, E. et al. 2007. Capturing the fugitive: applying remote sensing to terrestrial animal distribution and diversity. - Int. J. Appl. Earth. Obs. Geoinf. 9: 1–20.

28. Lillesand, T. M. et al. 2004. Remote sensing and image interpretation (5th ed.). John Wiley & Sons.

29. Madugundu, R. et al. 2008. Estimation of LAI and above-ground biomass in deciduous forests: Western Ghats of Karnataka, India. - Int. J. Appl. Earth Obs. Geoinf. 10: 211–219.

30. Mallows, C. L. 1973. Some comments on Cp. - Technometrics 15: 661–675.

31. Marasinghe, M.S.L.R.P. et al. 2015. Area suitability prediction for conserving elephants: An application of likelihood ratio prediction model. - Tropical Agricultural Research 25: 345–357.

32. Marshal, J. P. et al. 2010. Scale-dependent selection of greenness by African elephants in the Kruger-private reserve transboundary region, South Africa. - Eur. J. Wildl. Res. 57: 537–548.

33. McKay, G. M. 1973. Behavior and ecology: the Asiatic Elephant in southeastern Ceylon. -Smithson. Contrib. Zool. 125: 1–113.

34. Murwira, A. and Skidmore, A.K. 2005. The response of elephants to the spatial heterogeneity of vegetation in a Southern African agricultural landscape. - Landscape Ecol. 20: 217–234.

35. Myers, N. et al. 2000. Biodiversity hotspots for conservation priorities. - Nature 403: 853–858.

36. Ofstad, E.G. et al. 2016. Home ranges, habitat and body mass: simple correlates of home range size in ungulates. - Proc. R. Soc. B 283: 20161234.

37. Owen-Smith, N. and Chafota, J. 2012. Selective feeding by a megaherbivore, the African elephant (Loxodonta africana). - J. Mammal. 93: 698–705.

38. Owen-Smith, N. and Novellie, P. 1982. What should a clever ungulate eat? - Am. Nat. 119: 151–178.

39. Owen-Smith, R.N. 1988. Megaherbivores: The influence of very large body size on ecology. - Cambridge Univ. Press.

40. Pascal, J. P. 1982. Bioclimates of the Western Ghats. French Institute of Pondicherry, Pondicherry, Maps 1–2.

41. Pellew, R. A. 1984. The feeding ecology of a selective browser, the giraffe (Giraffa camelopardalis tippelskirchi). - J. Zool. 202: 57–81.

42. Pettorelli, N. et al. 2005. Using the satellite-derived Normalized Difference Vegetation Index (NDVI) to assess ecological effects of environmental change. - Trends. Ecol. Evol. 20: 503–510.

43. Pettorelli, N. et al. 2006. Using a proxy of plant productivity (NDVI) to track animal performance: the case of roe deer. - Oikos 112: 565–572.

44. Pettorelli, N. et al. 2011. The Normalized Difference Vegetation Index (NDVI): unforeseen successes in animal ecology. - Climate Res. 46: 15–27.

45. Prokopenko, C.M. et al. 2017. Extent-dependent habitat selection in a migratory large herbivore: road avoidance across scales. - Landscape Ecol. 32: 313–325.

46. Pyke, G. H. et al.1977. Optimal foraging: a selective review of theory and tests. - Q. Rev. Biol. 52: 137–154.

47. Rahman, D. A. et al. 2017. Factors affecting seasonal habitat use, and predicted range of two tropical deer in Indonesian rainforest. - Acta Oecol. 82: 41–51.

48. Reddy, C.S. et al. 2015. Nationwide classification of forest types of India using remote sensing and GIS. - Environ. Monit. Assess. 187: 1–30.

49. Rood, E. et al. 2010. Using presence-only modelling to predict Asian elephant habitat use in a tropical forest landscape: implications for conservation. - Divers. Distrib. 16: 975–984.

50. Rouse Jr, J. et al. 1974. Monitoring vegetation systems in the Great Plains with ERTS. - NASA special publication 351: p.309.

51. Roy, P.S. and Ravan, S.A. 1996. Biomass estimation using satellite remote sensing data—an investigation on possible approaches for natural forest. - J. Biosci. 21: 535–561.

52. Ryan, S.J. et al. 2012. The utility of normalized difference vegetation index for predicting African buffalo forage quality. -J. Wildl. Manage. 76: 1499–1508.

53. Saïd, S. 2005. Assessment of forage availability in ecological studies. - Eur. J. Wildlife Res. 51: 242–247.

54. Sannier, C.A.D. et al. 2002. Real-time monitoring of vegetation biomass with NOAA- AVHRR in Etosha National Park, Namibia, for fire risk assessment. - Int. J. Remote. Sens. 23: 71–89.

55. Sivaganesan, N. 1991. Ecology and conservation of elephants with special reference to habitat utilization in Mudumalai Wildlife Sanctuary, Tamil Nadu, South India. Ph.D. thesis. Bharathidasan Univ.

56. Sjöström, M. et al. 2009. Evaluation of satellite based indices for gross primary production estimates in a sparse savanna in the Sudan. - Biogeosciences 6: 129–138.

57. Srinivasaiah, N.M. et al. 2012. Usual populations, unusual individuals: insights into the behavior and management of Asian elephants in fragmented landscapes. - PloS One 7: p.e42571.

58. StatSoft, Inc. 2007. Statistica (data analysis software system), Version 8.0. www.statsoft.com.

59. Sukumar, R. 1990. Ecology of the Asian elephant in southern India. II. Feeding habits and crop raiding patterns. - J. Trop. Ecol. 6: 33–53.

60. Thakur, T.K. et al. 2017. Assessment of biomass and net primary productivity of a dry tropical forest using geospatial technology. - J. For. Res. pp.1–14 https://doi.org/10.1007/s11676–018–0607–8

61. Tsalyuk, M. et al. 2017. Improving the prediction of African savanna vegetation variables using time series of MODIS products. - ISPRS J. Photogramm. Remote Sens. 131: 77–91.

62. van Beest, F.M. et al. 2010. Forage quantity, quality and depletion as scale-dependent mechanisms driving habitat selection of a large browsing herbivore? - J. Anim. Ecol. 79:.910–922.

63. Varma, V. and Osuri, A.M. 2013. Black Spot: a platform for automated and rapid estimation of leaf area from scanned images. - Plant Ecol. 214: 1529–1534.

64. Wheeler, B.C. et al. 2013. Rates of agonism among female primates: a cross-taxon perspective. - Behav. Ecol. 24: 1369–1380.

65. Willems, E.P. et al. 2009. Remotely sensed productivity, regional home range selection, and local range use by an omnivorous primate. - Behav. Ecol. 20: 985–992.

66. Wilson, A.M. et al. 2011. Scaling up: linking field data and remote sensing with a hierarchical model. - Int. J. Geogr. Inf. Sci. 25: 509–521.

67. Wittemyer, G. et al. 2007. Breeding phenology in relation to NDVI variability in free-ranging African elephant. - Ecography 30: 42–50.

68. Wohlfart, C. et al. 2014. Mapping threatened dry deciduous dipterocarp forest in South-east Asia for conservation management. - Trop. Cons. Sci. 7: 597–613.

69. Wu, W. et al. 2013. Assessing woody biomass in African tropical savannahs by multiscale remote sensing. - Int. J. Remote. Sens. 34: 4525–4549.

70. Young, K.D. et al. 2009. The influence of increasing population size and vegetation productivity on elephant distribution in the Kruger National Park. - Austral Ecol. 34: 329–342.

71. Youngentob, K.N. et al. 2015. Where the wild things are: using remotely sensed forest productivity to assess arboreal marsupial species richness and abundance. - Divers. Distrib. 21: 977–990.

72. Zengeya, F.M. et al. 2013. Linking remotely sensed forage quality estimates from WorldView-2 multispectral data with cattle distribution in a savanna landscape. - Int. J. Appl. Earth. Obs. Geoinf. 21: 513–524.

73. Zinner, D. et al. 2001. Distribution and habitat associations of baboons (Papio hamadryas) in Central Eritrea. - Int. J. Primatol. 22: 397–413.

